# Spatially mediated interactions shape founder-cell fitness and community assembly in multi-species soil bacteria

**DOI:** 10.64898/2026.06.26.734772

**Authors:** Tania Miguel Trabajo, Isaline Guex, Helena Todorov, Xavier Richard, Christian Mazza, Jan Roelof van der Meer

**Author notes:** corresponding author Jan R. van der Meer, Department of Fundamental Microbiology University of Lausanne, 1015 Lausanne, Switzerland.

## Abstract

Microbial communities spontaneously colonize pristine environments, yet how species growth kinetics and individual cell variation shape community assembly remains poorly understood. Here, we use time-lapse microscopy imaging to track the division of individual founder cells in communities composed of up to seven soil bacterial isolates grown on nutrient surfaces. With cell lineage tracking, we quantify species-specific absolute biomass formation and growth kinetics from early growth through stationary phase. The reproductive success of individual founder cells depended on the timing of their first cell division, which determined their access to the primary substrate and their maximum growth rates. In mixed-species communities, founder cell success also depended on species-specific, substrate-dependent growth rates and yields. In addition, spatial factors – such as cells’ positioning, distances to non-kin neighbours, and identities of co-occurring species – further influenced outcomes. In spatially structured communities, interspecific interactions were globally governed by competition for primary substrates. We also observed cross-feeding of leaked metabolites, reflected in fluctuating paired interaction strengths and interaction signs. Species-pair interactions differed locally, with cells within distances of less than 15 µm exhibiting opposite interaction behaviours. Global pairwise interactions predicted from monoculture growth kinetics were observed in approximately half of the measured pairs, whereas measured paired interactions generally weakened in combinations of three or more species. Using a spatially explicit agent-based Monod growth model that includes interspecific interactions, we accurately predicted the compositions of seven-member communities. Overall, our results indicate that emergent, spatially mediated interspecific interactions between cells of different bacterial species primarily drive local and temporal changes in individual cell growth rates, which in turn determine final biomass formation. Because most natural microbial habitats are spatially structured, stochastic founder-cell positioning and fitness differences are key determinants of locally formed interaction patterns and species coexistence.

## INTRODUCTION

It is now generally accepted that interactions between cells of different species (i.e., interspecific interactions) play key roles in the spatial and compositional assembly of microbial communities and their functioning ^1–8^. Depending on their habitat, microbial founder cells will consume available substrates, divide and release metabolites that change the habitat’s chemical and nutrient composition. Differences in the inherent physiological capacities of individual taxa lead to varying degrees of their proliferation under the nutrient conditions prevailing in the habitat, influencing the taxa succession during the community’s assembly and its compositional outcome ^9–14^.

Interspecific interactions can take many forms but how these play out for a given habitat and community is poorly understood. Cells may kill or inhibit others through direct physical contacts by means of specific cellular appendages ^15, 16^ or through excreted small molecules^17^, enzymes^18, 19^ or released needle-like particles (e.g. tailocins^20^). Such microbial warfare processes have attracted wide interest because of their intricate molecular structures and mechanisms. The less sensational interactions occur as a result of the costless release of metabolites during microbial growth and their reciprocal utilization by other taxa that are present in the same habitat^3, 18, 21, 22^. The consequence of such metabolic cross-feeding is a loss of metabolic energy by the producer from consumption of primary (i.e., the initially available) nutrients, but potential gain of metabolic energy from consumption of secondary nutrients (e.g., metabolic intermediates) for secondary taxa unable to utilize the primary nutrients ^23^. Collectively, interspecific interactions are thus expected to influence community growth and taxa composition (i.e., the net sum of individual taxa population growth and decline over time), and to determine actual physiological activity and its associated metabolic reactions that drive broader-scale biogeochemical processes^7^.

Although the existence of the various interspecific interaction types has been well-documented, it is non-intuitive to describe how potential interactions are shaped in time and space, and quantify their importance for community assembly and behaviour^24–28^. Mostly, this is due to methodological difficulties to measure interactions, but also to the lack of accepted theoretical framework to link interspecific interaction parameters with community growth or decline, and extrapolate across scales of microbial activity in different habitats^23, 29–32^.

How are interspecific interactions measured? Microbial ecology is heavily leaning on sequencing methods to capture changes in taxa relative abundances between communities or over time^33–35^, or to infer system-wide changes in gene expression patterns of microbial taxa in specific habitats under environmental changes^36^. Such data can be used to deduce positively or negatively correlating taxa abundances over time or across habitats and build co-occurrence networks^33, 35, 37–40^. Laboratory experiments with defined mixed cultures can be used to infer interaction strengths from the observed differences in population growth in pairs or trios and monocultures^41^, and to predict growth outcomes in higher order mixtures from e.g., pairs (with varying degrees of success) ^42–44^. Testing growth effects for all possible taxa combinations even for small communities quickly becomes very cumbersome, which has spurred usage of computational predictions of nutrient-niche overlaps and growth outcomes from genome-scale metabolic models ^7, 41^. Improved microcultivation methods have also led to a massive scaling up of parallelized testing of growth effects of paired or higher order strain mixtures on fluorescently labelled focal strains^1, 2, 45^. In addition, interaction effects have been inferred from single strain dropout experiments in synthetic communities^4^, or by machine-learned trend predictions^25^. However, what has been missing so far are direct measurements of the dynamic nature of interspecific interactions and their effects on growth kinetics of individual community members. In addition, since most natural habitats for microbial growth are spatially structured, it is also pertinent to study how spatial positioning of cells shapes interaction-mediated effects. This combination could potentially widen classical microbial growth theory for single species ^46–48^ to community growth under inclusion of interspecific interactions and space^23, 49^.

The major objective of our study was thus to quantify growth kinetics and emerging interspecific interactions between bacterial species in a spatial context. We do this by direct time-lapse microscopy observations of growing populations from randomly positioned individual founder cells of monocultures, paired or higher order species mixtures (Fig. 1a)^50^. Cell division and formation of microcolonies occurs at rates dependent on their inherent metabolic capacities and controlled by diffusion of the available nutrients and electron acceptors. As a result of their metabolism, dividing cells deplete the nutrients but may leak or excrete metabolites, some of which can be used by themselves later on, or by neighbouring cells of other species (which may reciprocally do the same), thereby creating spatial, distance-and time-dependent interactions^51^. In the closed batch set-up of our experiments there is no new influx of nutrients and the different strains will go through all growth phases (i.e., lag, exponential, stationary). By reconstruction of spatially resolved species-specific cell lineages, differences in individual cell division rates of mono-versus cocultures become a proxy for both positive (i.e., cells growing better when being close) or negative interactions that can be summarized across many cell lineages across space. Finally, we deploy an agent-based cell model for microbial growth under carbon-limited conditions^49^ using Monod-type parametrization in a substrate-diffusible spatial context^50^ to test predictions of mono-or paired culture-derived growth and interaction parameter values (‘potential interactions’) for higher order assembly outcomes (‘realised interactions’).

**Figure 1.**
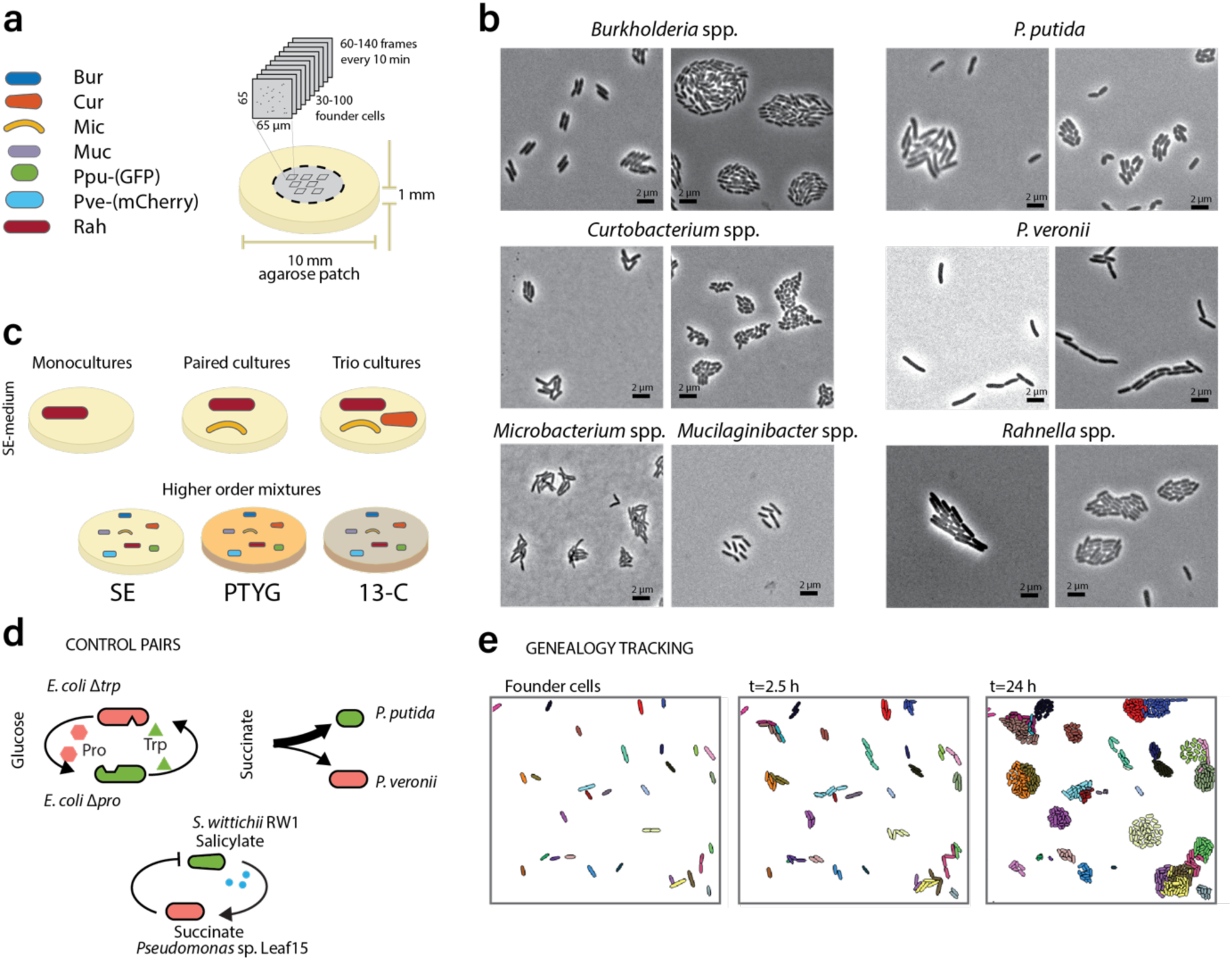
Experimental design and methodology. **a** Concept of random seeding individual founder cells on agarose patches. Seven isolates making up the soil mini synthetic community (Bur, *Burkholderia* sp.; Cur, *Curtobacterium* sp.; Mic, *Microbacterium* sp.; Muc, *Mucilaginibacter* sp.; Ppu, *Pseudomonas putida* – GFP tagged; Pve, *Pseudomonas veronii* – mCherry tagged; Rah, *Rahnella* sp.). Cell division into microcolonies is followed by time-lapse microscopy across predefined imaging areas, always situated in the patch centre. **b** Species identity of microcolonies is deduced from differences in fluorescence (Ppu, Pve) and characteristic morphologies during growth. Images show phase-contrast. **c** Strains are tested in monoculture, pairs, certain trios and full mini-SynComs. SE, soil extract; PTYG, peptone-tryptone-yeast extract-glucose medium; 13-C, 13 individual carbon substrate mixture. Carbon addition is limited to 1 mM or 100 mg l^-1^ to avoid multi-layered colony growth. **d** Additional strain pairs for benchmarking known interaction types (dependency, competition, inhibition). **e** Example cell tracking outcomes and genealogy/lineage building (illustrated in different colors).

## RESULTS

### Quantifying interspecific interactions from time-lapse microscopy of growing populations

We constructed a simplified soil microbial community (soil SynCom) of seven culturable soil isolates (a *Burkholderia* species, Bur; *Curtobacterium*, Cur; *Microbacterium*, Mic; *Mucilaginibacter*, Muc; *Pseudomonas putida*, Ppu; *Pseudomonas veronii*, Pve; and *Rahnella*, Rah) that are commonly found in natural soil communities^9^. Growth of the community was followed at single cell level by time-lapse microscopy across different strain order complexities, from monocultures to the complete seven strain coculture (Fig. 1). The soil isolates were chosen such that they could be reasonably well differentiated on microscopy images based on a combination of single cell and colony morphologies (Fig. 1b, *Methods*). Additionally, two of the strains (Ppu and Pve) were genetically tagged with low constitutive fluorescent protein expression to aid in species identification (Fig. 1b). Monocultures, different paired, trio or higher order combinations of soil isolates and the full mini-SynComs were surface inoculated, targeting between 50–150 randomly placed founder cells per imaged area and approximately equal founder cell numbers of each species in the mixtures (Fig. 1c). In addition, we tested three strain pairs with previously described interactions as controls for different interaction types (Fig. 1d). Mono-and mixed cultures were then incubated and imaged by time-lapse microscopy every 10 min until stationary phase was reached. Unique cell lineages (i.e., each from a unique founder cell in the earliest image) were reconstructed from cell segmentation, spatial positioning and cell tracking on consecutive time-lapse image series (Fig. 1e).

Importantly, individual cell data are inherently noisy and we use both single cell-derived measurements and averaged behaviour across imaged fields (FOI, or Field of Image) or multiple experiments to describe spatial-mediated interaction effects. Benchmarking of averaged behaviour of control strain pairs qualitatively recapitulated previously described interaction types (Fig. 2). For example, two auxotrophic strains of *E. coli*, one of which is unable to produce tryptophan (Δ*trpC*-RFP) and the other proline (Δ*proC*-GFP), positively influenced each other in coculture, characteristic for growth metabolic dependency^51, 52^. Biomass production of each of the strains in coculture with glucose as sole carbon and energy substrate was unaffected, but both strains grew on average 1.4-1.5 times faster in mixture than alone (Fig. 2a, *P* = 0.0021 for Δ*proC*, *P* = 0.1447 for Δ*trpC* in two-sided Wilcoxon ranksum test of position means). However, growth outcomes in terms of mean occupied cell area per FOI were opposite from the maximum specific observed growth rates of the monocultures (Fig. 2a, total area). The second pair consisted of a leaf isolate (*Pseudomonas* Leaf15) described to produce potentially growth-inhibiting compounds^53^; here tested in conjunction with *S. wittichii* RW1^54^ on a mixture of two selective carbon substrates, succinate for Leaf15 and salicylate for RW1. Paired incubation led to a statistically significant increase of Leaf15 growth rates and biomass formation, and decrease of RW1 growth, but an overall increase of the paired productivity sum (Fig. 2b). This is a sign for Leaf15 using leaked RW1 metabolites from salicylate to augment its own productivity. Example single cell data show global decrease of cell growth rates in early and late exponential phase, with considerable variations among individual cells. Colony sizes varied across space (Fig. 2c). One can also see how growth of *Pseudomonas* sp. Leaf15 is sustained for longer in coculture with *S. wittichii* with certain Leaf15 cell zones close to *S. wittichii* colonies having distinctly higher growth rates, suggestive for the overall observed average growth rate increase (Fig. 2b &c, arrows). The third benchmark used *P. putida* and *P. veronii*, as an example of competition for a single substrate, succinate (Fig. 2d) ^50^. Primary substrate competition here is evident from the sum of the paired productivities being equal to either mono-culture productivities, but *P. putida* producing a bigger share in the coculture than *P. veronii* (Fig. 2d). The reason for this is the faster growth rate of *P. putida* than that of *P. veronii*, both of which remain the same in mono-and coculture (Fig. 2d). Despite this global behaviour, single cell data show important growth rate disparities, leading to local variations of reproductive success (Fig. 2e). Overall, these results indicated that we could use single-cell spatial time-lapse measurements to characterize emerging interactions among the soil isolates.

**Figure 2.**
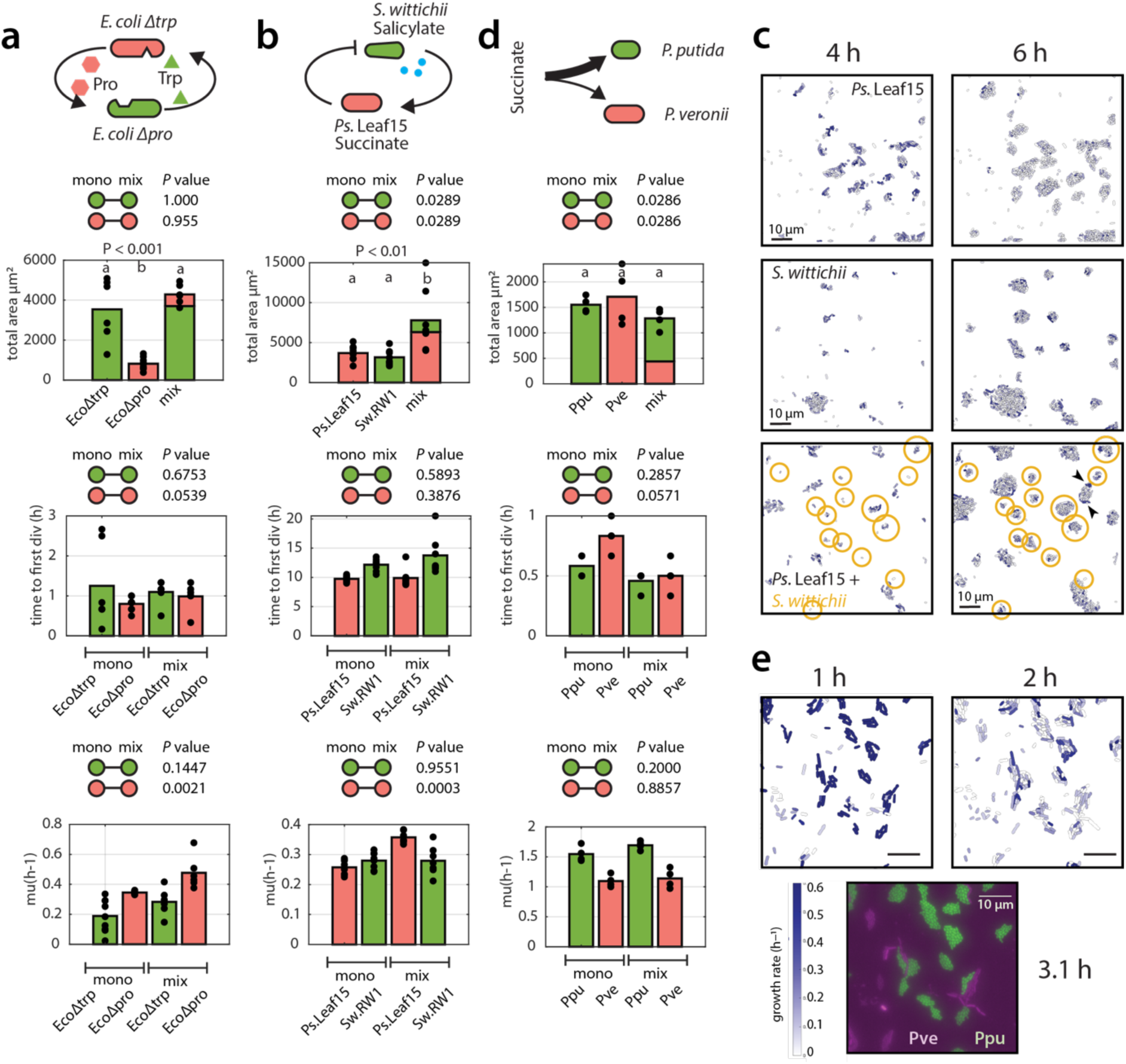
Benchmark outcomes to typify known interactions. **a** Mono-and coculture total occupied cell areas (in µm^2^ per image field), time to first division (in h) and maximum exponential lineage growth rates (mu, in h^-1^) for the dependency pair of *E. coli* Δ*trp* (cannot produce tryptophan) and *E. coli* Δ*pro* (no proline). Bars shown the mean of n=3-7 image field replicates (black dots). *P* values from two-sided Wilcoxon test between mono and coculture (mix) replicate values. (a, b) Grouping from one-way Anova and post-hoc testing. **b** As for (a) but for *Pseudomonas* sp. Leaf15 and *S. wittichii* RW1 growing on the combination of succinate (for Leaf15) and salicylate (for RW1). Blue dots indicate potential release of metabolites profitable for Leaf15. **c** Reconstructed field of imaging (FOI) with spatial cell positions from the segmented geometry measurements at mid and late exponential phase for mono-and cocultures of Leaf15 and RW1 (encircled in yellow). Colors indicate growth rates (scale in panel (e)). Arrows point to suggestive signs of increased Leaf15 cell growth rates nearby RW1. **d** As (a) but for the direct competition between *P. putida* and *P. veronii* growing on succinate as single primary substrate. **e** as (c) but for the coculture pair of d. Micrograph shows the specific fluorescence of Ppu (green) and Pve (mCherry) cells.

### Individual microcolony reproductive success is mostly determined by timing of first cell division

Next, we tested surface growth of the seven soil isolates in monocultures using the same agarose disk setup (Fig. 1a &b). For these experiments, we primarily used soil extract (SE) as mixed substrate, which may obscure certain dependencies developing under defined substrate-strain conditions (e.g., as for *Pseudomonas* Leaf15 and *S. wittichii* RW1 in Fig. 2b &c), but allowed consistent growth of all mini-SynCom members and can be considered representative for substrates found in soil^9^. Mean FOI-averaged monoculture growth characteristics of the 7 strains varied widely, with Bur displaying the highest productivity in terms of formed colony area, and Ppu and Pve the lowest (Fig. 3a). Ppu and Rah showed the highest, and Muc the lowest mean monoculture µ_max_ (Fig. 3b). Muc showed the longest average lag time (time to first cell division), and Bur-, Cur-and Rah-lag times were consistently among the shortest (Fig. 3c). Ppu, Pve and Rah were among the strains with the highest proportion of non-growing cells seeded on the surface (Fig. 3d).

**Figure 3.**
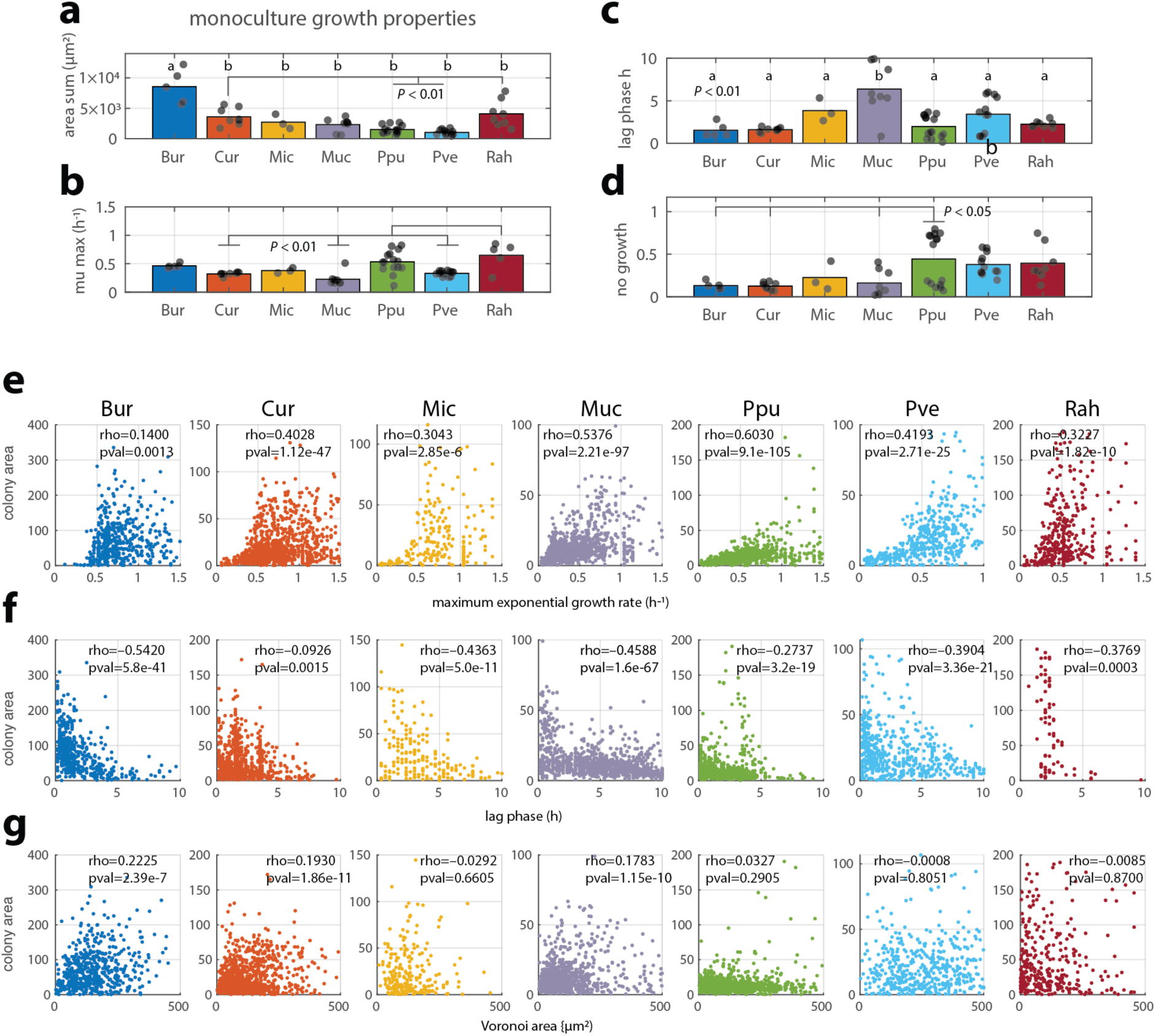
Growth kinetic parameters of the soil isolates in monoculture. **a** Total formed biomass area (in µm^2^ summed segmented cell area per field of image – FOI**. b** Mean maximum specific cell lineage exponential growth rate (µ_max_ in h^-1^, averaged over all cell lineages per FOI). **c** Mean time to first cell division (taken as lag time, in h). **d** Proportion of non-growing founder cells. Bars show means per FOI from all replicates (black dots). Letters signify significantly different grouped values from one-way Anova and post-hoc testing (highlighting individual group differences by indicated *P* values). **e** Microcolony area (in µm^2^) dependency on measured maximum exponential growth rate (h^-1^). **f** Correlation of time to first division – lag phase (h), and final microcolony/lineage area. **g** Correlation of Voronoi area around founder cell (in µm^2^) and final microcolony area. Dots indicate individual cell lineages. Rho and pval correspond to the calculated Pearson correlation coefficient and its *P* value, respectively.

In comparison to FOI-averaged kinetic parameters, individual microcolony properties vary widely (Fig. 3e-f; Supplementary figures 1 & 2). All monocultures showed a general positive and statistically significant correlation of maximum observed colony growth rates and attained stationary phase area (Fig. 3e). In contrast, stationary phase colony areas correlated negatively with the time until first division of the respective founder cell (Fig. 3f), and rather poorly to the available colonization space (Voronoi area, Fig. 3g). This suggests that reproductive success of founder cells relies heavily on their timing of first cell division, and less so on their positioning relative to others. Early dividers profit from available nutrients to develop faster growth rates, whereas late starters face an immediate nutrient shortage because of its consumption by others, slowing their growth rates (Fig. 3f). FOI-averaged growth characteristics thus compare more consistently than individual microcolony properties, which are influenced by stochastic lag time differences of their respective founder cells and their random starting positioning.

### Global interactions in among soil isolates are dominated by substrate competition

In order to determine the potential types and strengths of interspecific interactions, we next cocultured pairs, trios and higher order mixtures of the soil isolates on SE-agarose (Fig. 1a). Productivity changes in pairs compared to monocultures were in some cases reciprocal (i.e., one partner significantly higher, the other significantly lower than in monoculture, 1 out of 17 pairs, example Pve and Bur; Fig. 3a), in other cases both reduced (e.g., Bur and Ppu, 2 out of 17 pairs; Fig. 3a), negative on one partner (e.g., Mic and Muc, 9 of 17 pairs), or indifferent on both (5 of 17 pairs, Fig. 3a). In none of the pairs, the total biomass productivity was higher than either of its respective monocultures, whereas for four pairs the total biomass of the mixture was higher compared to one of the partners but no different compared to the other, or higher than one but lower on the other (2 pairs, Supplementary figure 3). In two cases (Bur + Ppu and Bur + Pve) the total biomass was significantly lower compared to one of the monocultures but higher to the other. For the remaining 8 pairs, the total biomass was indifferent compared to either of the monocultures alone, suggesting they are purely substrate competitive (Supplementary figure 3). Pairs with lower total biomass than expected from either partner are indicative for some inhibition other than competition for the substrate, which would result in the sum of the pair being no different than either monoculture (as in Fig. 2c). Also the FOI-mean maximum specific growth rates of some strains in pairs were significantly affected by the partner (Supplementary figure 4a). But there was no specific correlative trend between the ratio of the paired versus mono-culture growth rates (Supplementary figure 4a) and the corresponding productivities (Supplementary figure 4b, *P* = 0.2449, Pearson’s rho = 0.2050).

With few exceptions, the species decreased their productivity (mean FOI-summed cell lineage area) in higher order mixtures compared to mono culture (Fig. 4b), indicating that mini-SynCom growth and variations of lower order are competitive. Differences in FOI-averaged total productivity among all tested monocultures and mixtures on soil extract were significantly explained by the condition *trio* and presence of Bur (contributing positively to productivity), and negatively by Pve and Ppu (Fig. 4c). Notably Bur-productivity decreased in paired-cultures whereas Pve frequently gained in productivity, suggested by the mixed effect model (Fig. 4c). On the other hand, the total productivity in higher order mixtures as the FOI-summed areas of all microcolonies did not generally decrease compared to monocultures (Fig. 4c).

**Figure 4.**
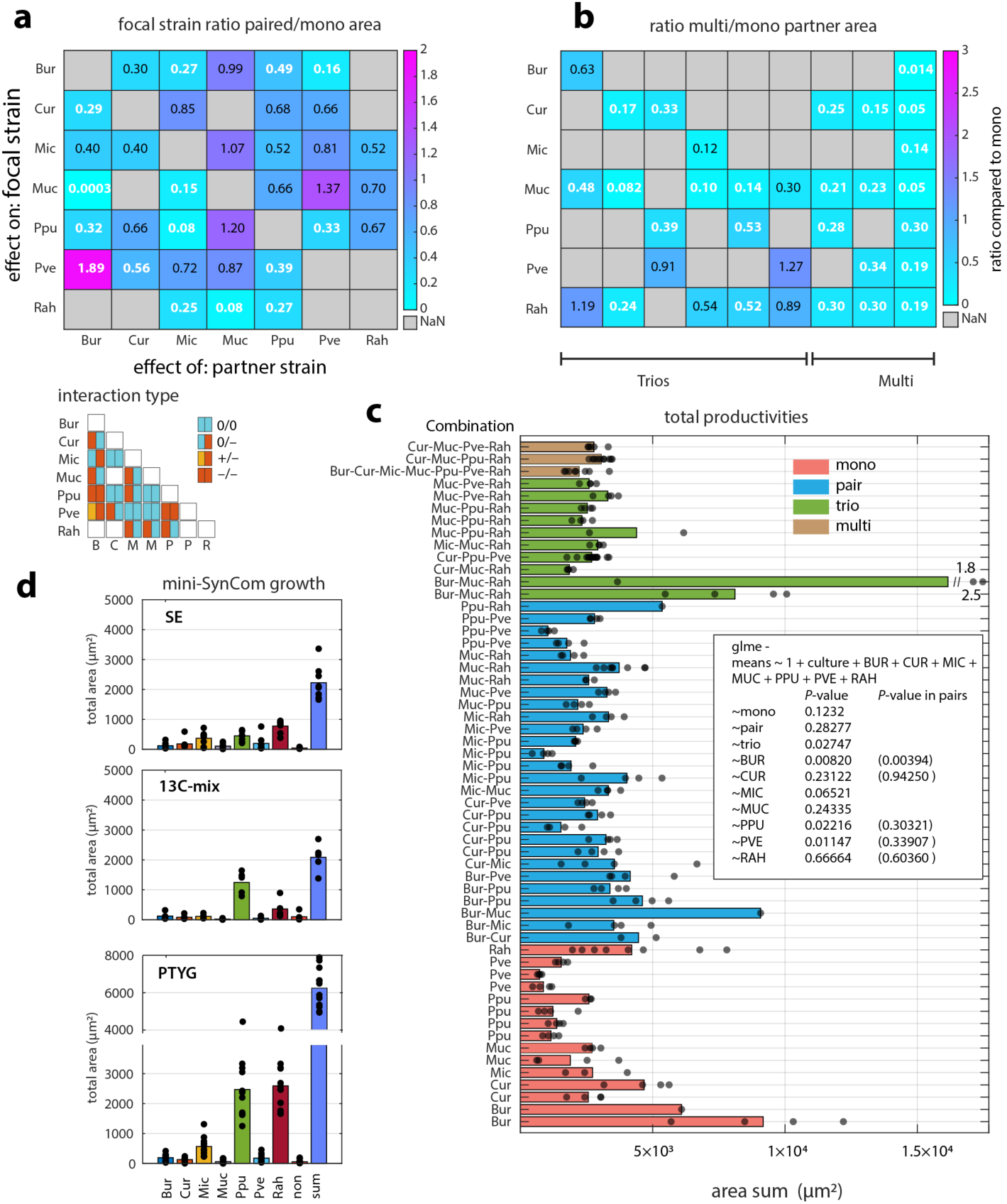
Global growth outcomes of the soil mini-SynCom members in pairs or higher order mixtures. **a** Ratio of mean total species biomass area per FOI in paired versus monoculture. Ratios shown as colour change according to scale on the side and with their mean values reproduced within each squared segment. Values in white bold type-face have a *P* value <0.05 in two-sided Wilcoxon test. Diagram below simplifies ratio changes as interaction signs (+,–,0; left color is focal strain, right is target strain). **b** as (**a**) but for ratios of higher order mixtures and monoculture productivity. **c** Mean total productivities (as summed segmented covered cell areas in µm^2^ per FOI) across all mono, coculture or higher order mixed culture experiments (compositions indicated on the left). Independent biological replicates kept separately. Bars show the mean of technical replicates (black dots, FOI-averages per experiment). Generalized linear mixed effect model (glme) tests effect on mean total productivities of culture type (e.g., mono, pair) or presence/absence of individual strains. **d** Mean per strain total productivity (FOI-averages, in µm^2^) in the mini-SynComs grown on SE (soil extract), PTYG or the 13 carbon compound mixture.

The final composition of the mini-SynCom on soil extract was dominated by Rah, Ppu and Mic (Fig. 4d). In contrast, with the mixture of 13 substrates the mini-SynCom was dominated Ppu, whereas with PTYG (peptone, tryptone, yeast, glucose medium, see *Methods*) Rah and Ppu dominated the community stronger (Fig. 4d). Notably, the maximum specific growth rates of Cur, Muc and Pve significantly increased in higher order mixtures (Supplementary figure 4c), whereas the trend between the strain’s ratios of growth rates and productivities in higher order mixtures versus in monoculture was weakly negative (influenced by the imbalance of datapoints, Supplementary figure 4d, rho = –0.4110, *P* = 0.0175). Overall, these data suggested that species interactions among soil isolates are competitive but there is no intuitive trend to extrapolate behaviour in higher order mixtures from global growth kinetics of mono-or paired cultures.

### Interaction strengths between species change over time and do not reflect direct substrate competition

To enable dynamic measurements of the strengths and directions of emerging interactions, we turned to a temporal analysis of local productivity differences between species. We previously found that a paired interaction can be described by the ratio of biomass productivity of either partner as a function of their founder cell ratio within a defined area^50^. This can be implemented by quantifying the cell ratios between ‘kin’ and non-kin taxa in a pair and the resulting summed biomass area ratio at any time point, within circles of increasing radius around every founder cell (Fig. 5a). During exponential growth this relationship is a function of the maximum growth rates of either species (Fig. 5a), and can be approximated by the slope from the linear regression of the ratio measurements (for either of the species, Fig. 5b; named here the competition index). When selecting only significant slope values (a Pearson correlation *P* value < 0.05 and linear regression coefficient > 0.2), the dynamic changes in paired interactions can be visualized (Fig. 5c). As an example for two independent experiments of the Bur + Ppu pair, in one case the slope of Bur versus Ppu is higher than 1 (Bur is the more competitive strain) but slowly decreases to below 1 (independent of the radius of the interaction circle), whereas from Ppu-perspective the slope starts lower than 1 but then increases to ca. 1.5 (Fig. 5c). In this case, the competition index of Ppu increases more in the smaller circle radii, suggesting that the closer it is to Bur the more it gains in competition strength over time, perhaps because Ppu profits from Bur-leaked metabolites. In a second independent experiment (Fig. 5c, Exp 2), the competition indices reverse and result in Bur producing more biomass (as colony area) than in Exp 1, and Ppu less. The reason for this different behaviour of the pair is the ca. 3-fold increased lag phase of Ppu in Exp 2 compared to Exp 1 (Fig. 5c), which gives Bur a growth head-start. This longer lag phase may be due to slight variations in the preculture preparation. An overview of the interaction dynamics between all strains and independent replicates of the paired experiments supports this general trend: interactions are not stable over time, change as a function of distance and can even vary in their sign among paired replicates because of stochastic differences in e.g., lag times (Supplementary figure 5).

**Figure 5.**
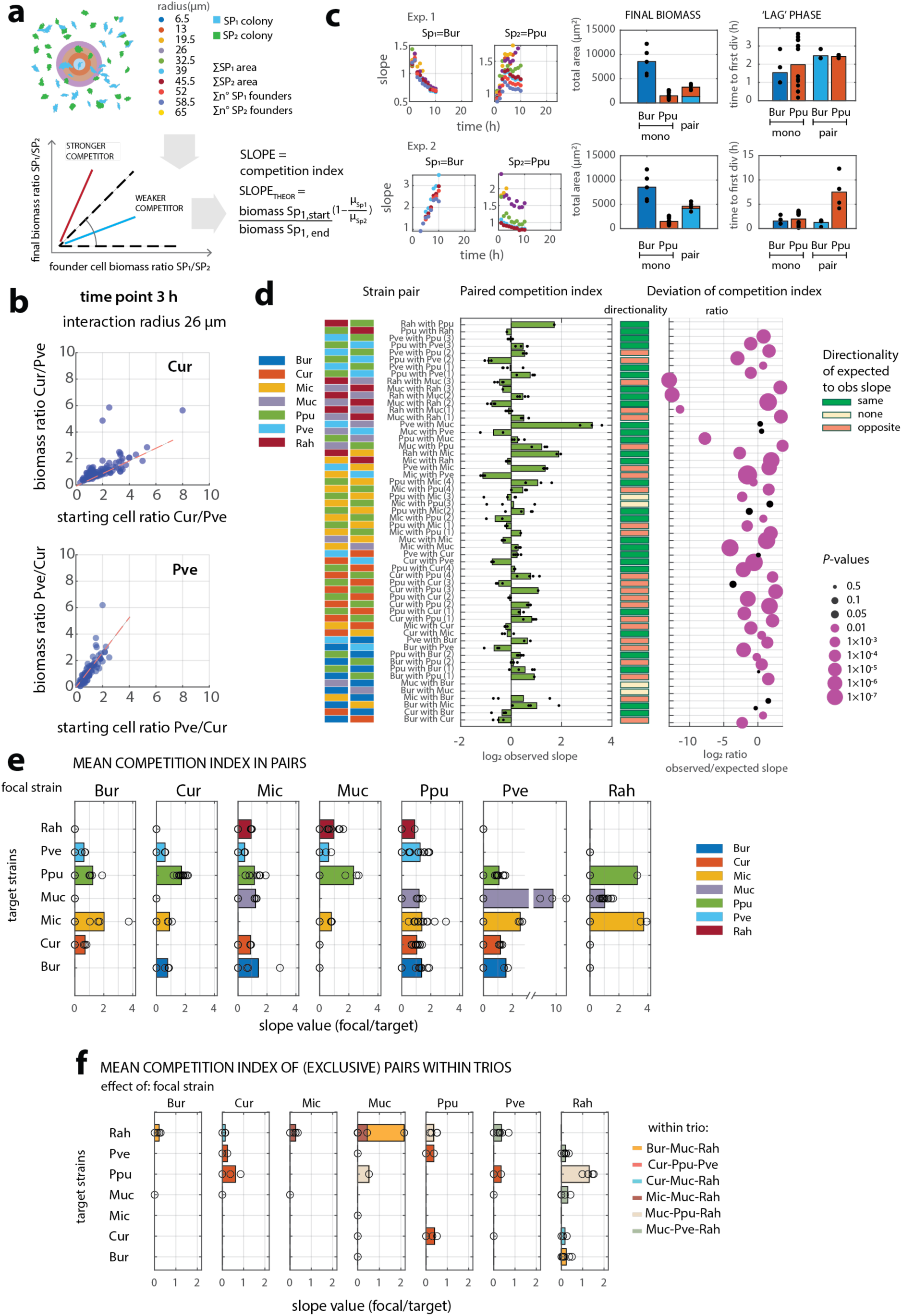
Population-averaged interactions among soil isolate pairs. **a** Principle of calculation of the paired competition index from the relation between the paired species founder cell ratios and their attained biomass ratios (as lineage areas) at any time point. Relation is measured at any time point for increasing circle areas placed on every growing founder cell (founder cells closer to the image boundaries than the circle diameters are excluded). This results in determination of slopes (**b**) that indicates the competitive strength of one paired member over the other (and reciprocally). **c** Slope values change dynamically during paired growth and can vary as a function of the circle diameter, pointing to closer or farther interaction ranges (dot colours correspond to circle diameter scale of panel **a**). The onset and direction of the slope development can depend on the lag time of founder cells (as shown here for two independent biological replicates of the Bur+Ppu coculture). Only slopes with Pearson correlation coefficient < 0.05 and r^2^ linear regression < 0.2 taken into account. Full data set in Supplementary figure 5. **d** Observed mean competition indices across all independent biological paired replicate experiments (composition shown on the left, colours corresponding to species colour scale) in reciprocal species order per pair. Bars show the means of technical replicates (black dots, representing the FOI-average from slope values at exponential phase time points (1–6 h) and for circle diameters between 19.5–45.5 µm. Direction (+, – or 0, in colours) and deviation of observed competition indices to the calculated slope based on measured monoculture growth rates (e.g., from Fig. 4a; formula in panel **a**). Circle diameter of points corresponds to *P* values from two-sided t-testing of calculated and observed replicates. **e** Mean competitive index per focal strain with the indicated target (bars show the average across different biological and technical replicates shown as open circles; values from panel **d**). Bars higher than 1 indicate the focal strain is more competitive than the target. **f** as for (**e**) but for paired slope interactions measured within trios (focusing on circle areas exclusively occupied by only 2 of the 3 members).

To summarize the comparison of the different pairs and across experimental replicates, we averaged the slopes during exponential phase and from three interaction circle radii (19.5 – 39 µm), for which the Pearson and linear regression coefficients were the most frequently statistically significant (Fig. 5d, Supplementary figure 5). This averaging (Fig. 5e) indicates, for example, that Bur is the more competitive in pairs with Ppu or Mic, but less so with Cur or Pve (slopes smaller than 1). Cur, Muc and Rah are also more competitive than Ppu (slopes higher than 1), whereas Ppu, Pve and Rah are more competitive than Mic (Fig. 5e). For several pairs, slopes and slope differences oscillate closely around 1 indicating neutral interactions, and are not always reciprocal (likely because of the simplification of the averaging in Fig. 5e, see Supplementary figure 5b). The measured average paired slopes can be compared with slope values predicted from the monoculture growth rates, according to the theoretical relationship presented in Fig. 5a, which would indicate direct substrate competition. Predicted competition indices calculated from monocultures have the same directionality as experimentally observed in ca. half of the pairs, whereas they are opposite or not visible (competition index close to 1) for the other half (Fig. 5d). Quantitatively, the predicted competition index is 2-4 fold and for most pairs significantly different than what is actually measured (Fig. 5d). This indicates changes in growth kinetics in pairs with respect to either monoculture that go beyond direct substrate competition and point to cross-feeding or inhibitory metabolite effects.

We were also curious to understand how the competition indices change in higher order mixtures. However, even though one can conduct the same type of analysis as in Fig. 5a, there is no theoretical basis for comparison of the slope lines from individual monoculture growth rates in higher order mixtures. The only possibility is to detect local exclusive pairs within trio (or higher order) mixtures, focusing on areas with only two out of three (or more) species being present within the circle ranges used for the analysis (Fig. 5a). Among the trio mixtures tested, we noticed a tendency from slope measurements that exclusive pairs in trios lose competitiveness compared to paired cocultures (Fig. 5f). For example, the Cur-Ppu pair within the Cur-Ppu-Pve trio has a slope lower than 1, whereas it was higher than 1 in Cur-Ppu paired experiments (Fig. 5f). The only exception we found here was the higher competitiveness of Muc with Rah in the trio with Bur (Fig. 5f). Measurements of paired slopes within mixtures of more than three species were inconclusive because of the scarcity of spatial areas with sufficient exclusive species pairs to calculate slope values. This will be revisited by single cell measurements in the following section. These results thus showed that actual realised interactions in trio mixtures also deviate from interaction generalisations of paired measurements, and further suggest that emerging interactions in pairs and trios must go beyond simple competitive growth, in which case monoculture kinetic differences would be sufficiently explanatory.

### Local changes in cell division rates show time-, distance-and strain-dependent interactions

To better characterize distance-and time-dependent interactions between strain pairs even in higher order mixtures, we focused on changes in individual cell division rates over time as a function of distance. As mentioned for the paired-strain benchmarking (Fig. 2c & e), this comes with an increase in variability and noise of measured cell behaviour, and potentially more difficulty to substantiate significant effects. Hence, we ask the question whether cell division rates are influenced by the closest neighbour species identity. We differentiated here whether cells were exclusively surrounded by their own kin or by cells from non-kin species (as illustrated in Fig. 6a), scoring the proportion of cells with statistically significantly deviating cell division rates nearby a non-kin cell compared to those of exclusively their own kin at the same time point. For every cell we focused on areas of increasing radius up 15 µm, scoring exclusive own kin areas as those where no neighbours of other strains were present (Supplementary fig. 6a). Cell division measurements in exclusive ‘own-kin’ areas within growing mini-SynComs showed similar mean maximum growth rates as in monocultures except for Muc (Supplementary fig. 6b, *solo* growth rates). In contrast, cells neighbouring different species frequently deviated from solo growth rates observed at the same time point. For example, the growth rate in a Cur-Ppu coculture of Cur-cells nearby Ppu first increases and then decreases, in comparison to Cur-cells that have no Ppu partners (Fig. 6b). Translated into proportions, more Cur-cells nearby Ppu have increased growth rates in early time points and decreased growth rates at later time points, whereas this is more or less opposite from the perspective of Ppu-cells (Fig. 6b). As suggested previously from the competition index measurements (Supplementary figure 5), independently repeated paired experiments can have different dynamics of growth rate changes because of variation in e.g., lag times of individual cells (Fig. 3c). In two out of four replicate experiments, Ppu-cells nearby Mic showed increasing growth rates, but in the other two replicates a more negative impact of Mic on Ppu is visible (Fig. 6c, Supplementary figure 7). In addition to alternating interactions signs over time, we also find evidence for simultaneous appearance of both positive and negative influence on growth rates of part of the population. As example, in the paired interaction of Ppu and Cur, about the same proportion of Ppu cells experience an increase as a decrease in local growth rates (Fig. 6d). This may be the result of differences in founder cell phenotypes and their lag times that influence the locally developing interaction.

**Figure 6.**
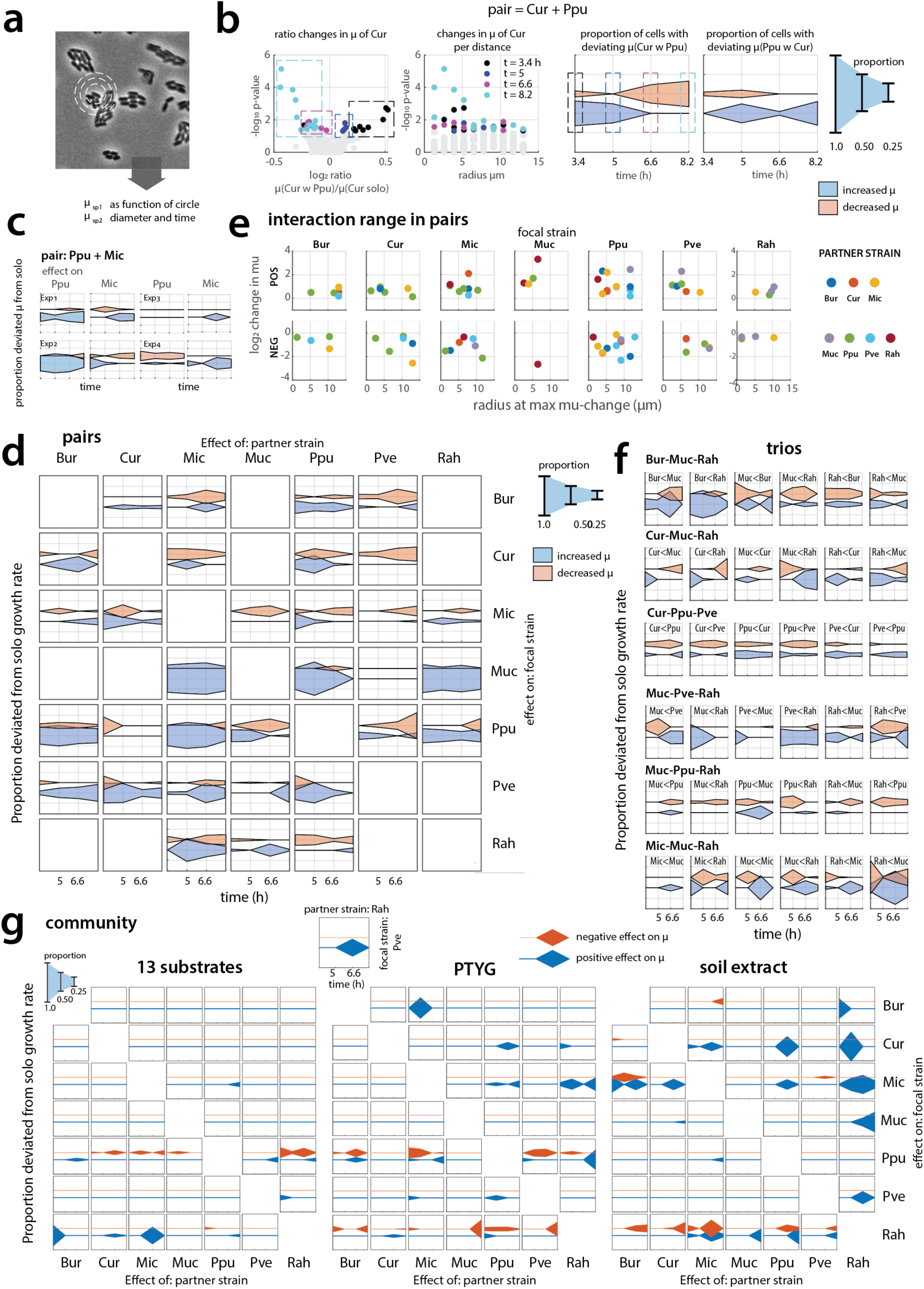
Dynamic paired-interactions in pairs or higher order mixed cultures of the soil isolates deduced from single cell neighbourhood measurements. **a** Individual cell division times are measured and averaged across circles of increasing diameter around each cell at defined time points to derive (**b**) the relative changes in growth rates of focal species nearby their paired target species, in comparison to growth rates of the focal species (in the same imaged area) exclusively surrounded by its kin. Statistically significant growth rate changes in presence of target species compared to kin only (Wilcoxon two-sided testing of individual cell values within the respective circle diameter, *P* < 0.05) are identified and score per time point (black, blue, magenta and cyan dots) and radius (rectangular dotted areas), and the proportion of cells with significant change compared to all cells in contact is plotted as a function of time in a reciprocal focal strain manner (e.g., Cur in presence of Ppu, or Ppu in presence of Cur). **c** Dynamic differences in paired interactions across multiple independent experiments of here Ppu and Mic (full overview in Supplementary figure 5). **d** Overview of interaction dynamics in all pairs. **e** Deduced interaction ranges for all paired measurements (from the maximum observed *P-*value change across radius distance, as shown in panel b). **f** Measured paired interactions in trio cultures. Indication ‘Bur<Mic’ meaning Bur is the focal strain (whose µ-change is depicted) under influence of Mic as the paired target strain. Proportions are here in comparison to all cells in contact with that partner, but not any other (region-exclusive). **g** Dynamic interactions measured for all (region-exclusive) strain pairs in the incubated mini-SynComs on the three different carbon source mixtures; cell pairs combined from all replicates (*n* = 8-10). Inset cartoon explains how to read the paired interaction in a single square from focal and target strain.

Across all measured pairs, Muc, Ppu and Pve showed the highest fractions of cells with significantly increased growth rates, suggesting they can profit from metabolites leaked from neighbouring paired non-kin cells, whereas the others show both positive and negative impacts varying over time (Fig. 6d, Supplementary figure 7 for a full overview of all pairs). The higher resolution single cell data support the measured differences in paired averaged compared to monoculture growth rates (FOI-averaged across lineages and all experiments; Supplementary figure 4). This indicates that coculturing more generally affects growth rates, and likely through the changed growth rates the final biomass ratios (Fig. 4c). The interaction ranges, taken from the position of the lowest *P* value for the µ-ratio comparison as a function of the analysis radius (as in Fig. 6b and Supplementary figure 7), oscillate between 1-15 µm, without clear partner-dependent influence and similar for negative and positive influences on the growth rate (Fig. 6e, Supplementary figure 7). Dynamic strain-pair patterns are retained in similar form within the tested trio combinations (for example, the more negative impact of Cur on Ppu and Pve, or the dynamically changing interactions signs of Ppu and Pve; Fig. 6d and f). In other cases, the presence of a third partner is changing the paired interaction; for example, Muc and Rah within the various trios (Fig. 6f and as suggested by the competition index slope measurements of Fig. 5f).

By quantifying growth rate changes in neighbouring cells exclusively limited to circle areas with only two species, also the effects of ‘pairs’ within the 7-membered mini-SynCom can be visualized (Fig. 6g). Mini-SynComs grown on SE become dominated by Rah (Fig. 4d), but the single cell dynamic growth rate analysis shows that all strains except Ppu profit during some period from being close to Rah-cells and showing faster growth (Fig. 6g). Reciprocally, Rah-cells show signs of being negatively impacted in their growth rate by most partners at some point during growth (Fig. 6g). Also Mic and Cur cells located nearby to Ppu showed temporary increased growth rates (Fig. 6g). Other paired interaction effects within higher order mixtures were not detected because of insufficient numbers of cell pairs in close distance (on average 23% of cells per species being engaged in exclusive pairs; Supplementary table 1 and 2). Mini-SynComs grown on the two other substrate mixtures (PTYG or the mixture of 13 different carbon substrates) displayed changed interaction patterns. For example, the driving role of Rah on SE was no longer visible, but instead more negative impacts on Ppu became visible, which had an overall more dominant role in those communities (Fig. 6g).

In summary, this analysis indicated that numerous dynamic interactions influence growth rates of neighbouring cells in a pair-dependent manner. These results further indicate that fast-growing dominant members in the community (like Rah and Ppu) exert a positive effect on others in form of temporally increased growth rates of nearby-located cells.

### Community growth depends on individual kinetic properties

As our results clearly pointed to local time-and distance-dependent dynamic changes and strong effects of stochastic founder cell variations (Fig. 2), we wondered as to how well the community composition outcomes can be explained from global growth kinetic properties of the individual members. To simulate community growth, we took advantage of a previously developed agent-based surface growth model^49, 50^ that was extended to 7 species. This model takes into account the experimentally measured starting positions of each cell and species, and calculates the resource utilization from diffusion and uptake conversion into biomass, with cells dividing and repositioning into microcolonies in the same manner as in the experiments. We then simulated four scenarios. In the first, cells were given the measured species growth rates and yields from the monocultures, plus the measured variation and the lag time information, as in Fig. 3. This would be equivalent to a scenario of competitive growth for substrate but no other interactions. In the second we used the monoculture measured mean per-species maximum growth rates but corrected by the observed change in paired-growth rates (i.e., values of Supplementary figure 4a) for founder cells that were nearer than 15 µm. In the third, we corrected the monoculture yields by the observed mean competition indices as in Fig. 5, again only for founder cells closer than 15 µm apart. In the fourth simulation we used the measured growth rates in the mini-SynCom with SE medium taken as the mean of the maximum observed exclusive single cell solo growth rates (Supplementary fig. 6b), while maintaining the respective monoculture yields (Fig. 3a).

Simulations with the measured mini-SynCom growth rates gave the most accurate results in the timing of community growth, compared to the experimental observations, whereas the other simulations predicted faster growth (Fig. 7a). All simulations broadly explained the total species formed biomass areas, with again the simulation with the measured mini-SynCom solo growth rates being the most correct, except for underestimating growth of Mic (Fig. 7b-e). Normalized relative abundances in all simulations except the one with growth-rate-corrections were similar, and not completely overlapping with the actual data (Fig. 7f). Finally, no common weight factor could be calculated by element-wise, linear least square, or quadratic fitting that would explain the difference in monoculture and mini-SynCom species growth rates from their relative founder cells proportions in the mixture, nor as an *a posteriori*-imposed criterium on the total community biomass from average competitive indices determined in paired cocultures (Supplementary data 1, Fig. 7e ‘pair-wise ratio constraint’). Even though monoculture-measured species growth rates gave reasonable prediction of mini-SynCom outcomes (Fig. 7b), the effective growth rates of founder cells in the mini-SynCom are a non-linear function of relative positioning, lag phase, interaction range to neighbouring cells, dynamic primary substrate utilization, and production and uptake of formed shared metabolites.

**Figure 7.**
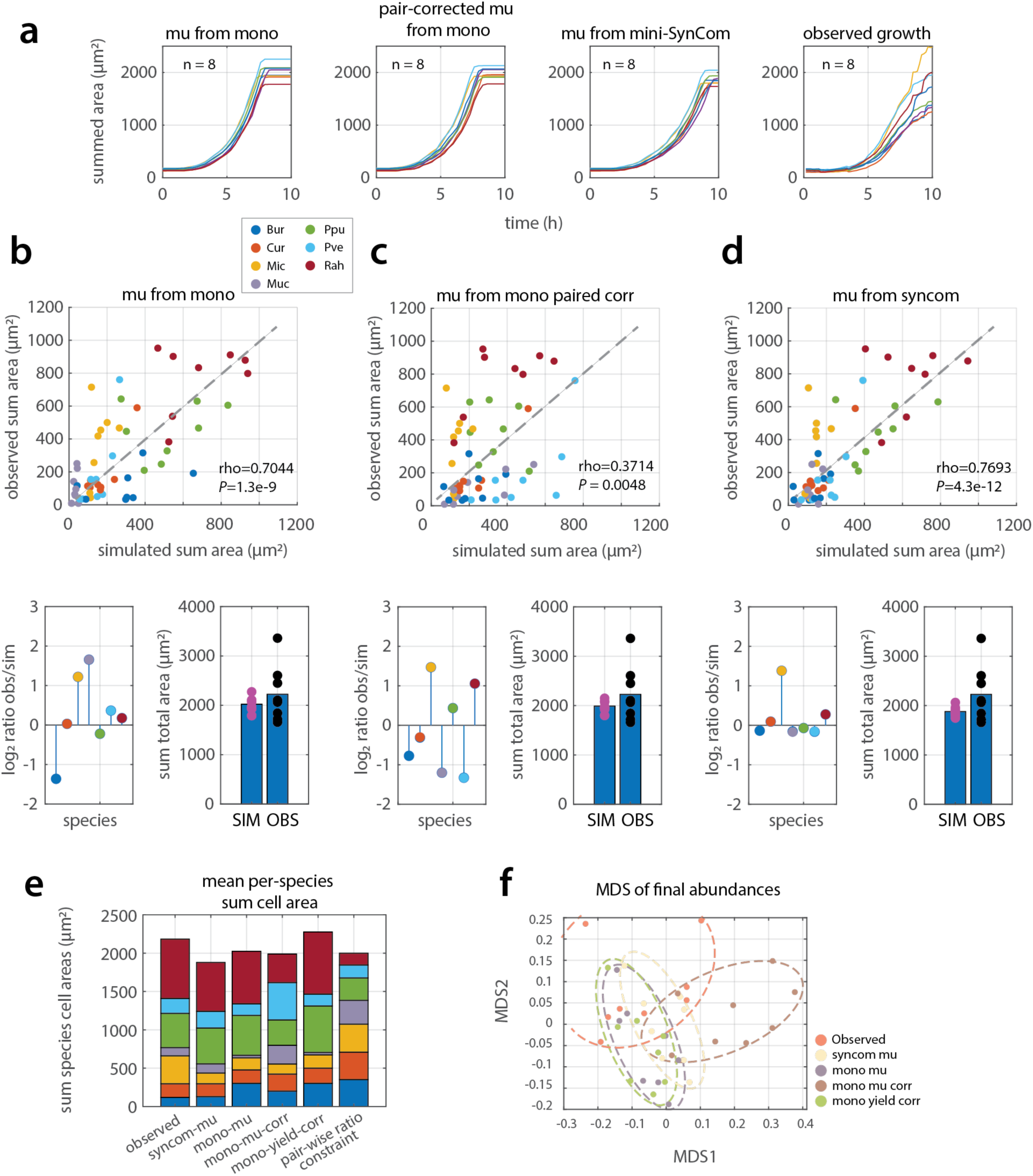
Simulated versus observed productivity outcomes of surface-inoculated mini-SynCom growth with soil extract. **a** Summed community growth (i.e., summed from all individual species lineages per image area, *n =* 8 replicates), using founder cell numbers and positions as in the experiments. Mu from mono, simulations seeded with maximum species specific growth rates, yields and lag times measured in monocultures from Fig. 3a-c. Pair-corrected mu from mono, as ‘mu from mono’, but with a correction on mu according to the measured paired change (as in Supplementary fig. 4) when founder cells of different species are within 15 µm. Mu from mini-SynCom, simulations seeded with mean species growth rates measured from kin-only single cells during exponential phase (‘solo’ growth rates; as in Supplementary figure 5). obs, experimentally observed community growth. **b - d** Individual summed species biomass per replicate for the simulations of (a) versus experimentally observed values (*P* and *rho* from Pearson correlations), their mean log_2_ ratio difference and the total sum comparison. Simulations calculate in formed biomass as a function of substrate consumption, which is then converted back to cell area using mean measured per species cell lengths and widths. **e** Absolute stacked per species abundances for the different simulations and observations. Mono-yield correction, corrected yields according as in Fig. 4a when paired founder cells are < 15 µm apart; pair-wise ratio constraint, estimation of the species abundances from the measured mean paired slope factors and the known starting proportions. **f** Multidimensional scaling of Bray-Curtis distances between normalized relative species abundances in the observed and simulated communities of (e).

## DISCUSSION

By conducting microscale surface growth experiments with monocultures, pairs, trios and higher-order mixtures (up to seven strains), we captured real-time changes in cell-division and lineage-growth rates as a function of spatial distance, founder cell positioning, species composition, and from early growth through stationary phase. Differences in FOI-averaged lineage-growth rates between monocultures and the same species in pairs or higher-order mixtures indicate species-and order-dependent interactions (Fig. 4 &5). Variations in cell division rates depending on neighbour distance and on primary substrate type directly demonstrate the dynamic nature of emergent interspecific interactions even in the full seven-member community (Fig. 6). Our results indicate that interactions are not constant over time, and, in a spatial context, are not a priori predictable for defined species pairs (Fig. 5). Interaction signs within pairs can alternate over time and opposite interaction signs can simultaneously coexist locally (Fig. 6). As a consequence of stochastic behaviour of founder cells, notably in the onset of cell division, interaction signs and outcomes can invert.

Paired-versus-monoculture growth outcomes have frequently been used to project community behaviour in higher species complexity ^1, 2, 42, 44, 45, 55–59^. Species pairing is experimentally straightforward, can be massively parallelized to include a large variety of nutrient conditions (e.g., Refs. ^1, 2, 45^), and allows classification of an interaction’s directionality and strength based on the paired growth outcome. Growth outcomes (or “yields”, definition depending on the method) are then combined with ad hoc models ^1^ or machine-learned training to fit factors that best explain higher-order community compositions from paired and/or trio mixtures ^42, 44, 55^. Although this has worked well in some cases (and less well in others ^43^), the ad hoc models do not connect to a general underlying growth theory. We agree that paired interactions can be seen as potential interactions for in mixtures of multiple species mixtures, but we also observed that paired interactions measured in pairs generally become weaker in trio settings (Fig. 5e &f), with some commonalities across pairs, trios and in the seven-member community (e.g., Cur-Ppu-Pve in Fig. 6f), but with mostly different actual interactions. Paired measurements (in pairs) thus tend to overestimate the magnitude of the realised paired interaction in higher-order species mixtures. This may explain previous observations that interactions become less ‘strong’ in higher-order mixtures than expected from pairs ^43^.

Instead of relying on extrapolations from pairs or trios to higher order interactions, we propose to include interspecific interactions in community growth models as a kinetic parameter in their own right, which acts by modifying cell growth rates. In a spatial context it might be sufficient to solely include paired interactions, since local areas rarely have cells of more than 2–3 species. The potential for paired interactions can then kinetically be described by the slope of the attained versus the initial species biomass ratio within defined spatial areas ^50^, defining the potential range of realised interactions. This would be similar as maximum growth rate and the half-saturation constant K_S_ describing the potential range of growth rates under given substrate conditions. For now we embodied this idea through an agent-based model with species founder cells initiated with their monoculture Monod kinetic parameters and their variability (maximum growth rates, yields, lag phases), adjusted for cells falling within short-range species interactions (i.e., closer than 15 µm). This is not completely elegant but showed that community growth kinetics, composition outcomes, and individual species abundances can be explained from first kinetic principles under inclusion of interspecific interactions (Fig. 7). Our results and models thus support previous modeling conclusions^30^ that the framework of classical growth kinetics is sufficient to explain and predict growth outcomes without invoking the signs, types and directions of species interactions. Improving community models along these lines could help to better predict community compositional changes and growth in various habitats based on monoculture growth kinetic and interaction measurements, or in future, based on modelled growth rates and potential interactions from genome-scale metabolic models ^7, 41, 60^.

Maintaining nomenclature of paired interactions (e.g., mutualism, ammensalism) can make sense in ecological context, but they may be the outcome of differences in monoculture kinetics without any additional species-specific interactions. Indeed, approximately half of measured averaged interaction pairs among the soil bacteria and the applied substrate conditions would qualify as being the consequence of substrate-dependent monoculture growth kinetic differences (Fig. 5d). However, this does not mean that the community develops without any additional emerging interactions: those not explained by substrate-dependent monoculture growth kinetic differences. More detailed real-time measurements of growing species-pairs indicate that interaction values change over time, whereas substrate competition based on the monoculture kinetics of either partner would result in a single value. In addition, we see that the competition index values change as a function of measured interaction range (Fig. 5, Supplementary figure 5), which must mean that growth rates vary locally and dynamically, depending on founder cell properties and their daughters (e.g., Fig. 2c &e); something that we confirmed by measuring division rates of individual cells within cell lineages and across independent experiments (Fig. 6).

At the used founder cell densities and substrate conditions, we find between 5 and 50% of cells in paired context with another species (i.e., within radial distances of 15 µm) display statistically significant growth rate variations compared to cells of the same species that are further apart (Fig. 6, Supplementary tables 1 and 2). These proportions were similar for measurements in pairs or higher order communities, but not necessarily showing the exact same timing or interaction sign, and simultaneous opposite interactions may appear within the larger FOI (Fig. 6). This is of course a consequence of the spatial setting and random placement of founder cells, which with increasing species complexity makes it increasingly unlikely that all potential paired interactions will emerge. Additional stochastic events in the onset of growth may further bias the actual growth of individual species into lineages that determine whether and which paired contacts will prevail. The combination of these factors contributes to some paired interactions dominating the community growth and others remaining without effect (as in Fig. 6g).

The advantage of our study setup is that it permits to extract both FOI-averaged growth kinetic paired behaviour as well as local variations that are the result of stochastic processes and cell phenotypic differences in a spatial context. We acknowledge that extracting local and temporal cell kinetic data is influenced by noise arising from technical difficulties of cell segmentation and tracking, but local patterns can be consolidated by taking many individual cell observations. One of the most important kinetic parameters determining reproductive success is the time until first cell division and less so the available space around a founder cell (Fig. 3). Faster start enables to develop maximum growth rates, because of the higher flux of available substrate under the initial conditions. Later starters find locally lower substrate fluxes, reducing their growth rate. Their growth may be penalized further by the faster-than-average primary substrate depletion from expanding colonies that bias substrate diffusion toward them as a result of faster uptake and locally lower concentrations. On the other hand, locally increased growth rates are hard to explain by local variations in primary substrate availability, and are more likely the result of local production of metabolites, formed by one species and preferentially used by others nearby. This concept of metabolite leakiness and reutilization has been demonstrated numerous times^3, 18, 21, 22, 61^, and is particularly relevant for a spatial setting where physiologically active cells become point sources for de novo diffusion of leaked metabolites that others can take up; the types of short-range interactions appearing in densely growing species pairs^51^. Variations of cell-division rates are thus likely a consequence of local and temporal changes in available substrates that originate from physiology and metabolism of neighbouring species. If its founder cells start dividing sooner, a cell lineage from a particular species may gain an advantage over a later-starting neighbouring species lineage, but this lineage may then profit from the faster-growing neighbour colony by uptaking leaked metabolites, as demonstrated in the support role of *Rahnella* (Rah) in the seven-member community growing on soil extract (Fig. 6g).

Our experiments and results of single cell and lineage growth changes give a different perspective on the roles and influence of emerging interspecific interactions in a community context. So far, there have not been any direct measurements of emerging interaction parameters in communities through growth rates, except for two cases of a single defined species pair^50, 62^. Instead, interactions are mostly deduced from differences in e.g., compositional changes or growth outcomes in a particular experimental or habitat setting, which tend to act as global population measurements summarizing the ecological success of multiple founder cells. For growth assays in bulk liquid suspension individual founder cell fates play a minor effect, because stochastic differences in the physiology of founder cells are lost as the populations become dominated by the fittest growers. However, for colonization of environmental or host habitats where spatial fragmentation may restrict mixing or where ecological bottlenecks restrict colonization to small-size founder populations, stochastic differences in viability and physiological status of single founder cells may become crucial for their reproductive success within communities. We would expect this to be the case for habitats such as skin^63^, micro-droplets^54^, particles^10, 12, 62^, plant leaves^4, 64^ or soil pores^65^. Local escape from or overturning of interaction signs as results of positioning and founder cell phenotypic heterogeneity may thus form important processes facilitating coexistence of slower-growing species amidst more competitive faster opportunists^50^.

## MATERIALS AND METHODS

### Bacterial strains and cultures

The bacterial strains used for the simplified community (mini-SynCom) included *P. putida* F1-eGFP (Ppu, strain 5790) ^50^, *P. veronii* 1YdBTEX2-mCherry (Pve, strain 3433) ^50^, *Burkholderia* sp. strain 6699 (Bur), *Microbacterium* sp. strain 6150 (Mic), *Curtobacterium* sp. strain 6155 (Cur), *Mucilaginibacter ginsenosidivorax* sp. strain 6152 (Muc) and *Rahnella* sp. strain 6700 (Rah) ^9^. Further strains used as examples for defined interactions included an eGFP-tagged strain of *Sphingomonas wittichii* RW1 ^66^ (strain 3354) and mScarlet-I-labeled *Pseudomonas* sp. Leaf15 ^53^, strain 7803 ^54^, *E. coli ΔproC*-GFP and *E. coli ΔtrpC*-RFP ^51^.

Pure cultures were maintained at –80 °C in 15% *v/v* glycerol and freshly regrown on nutrient agar plates to obtain individual colonies. For labeled strains, the antibiotic needed for selection of the genetic construction was added to the growth medium. For relatively fast growing strains (Ppu, Pve, RW1, Leaf15, *E. coli*), a single colony was transferred to liquid medium for preculturing, which was grown on R2A-medium for 24 h. For slow-growing strains we used nutrient-agar surface-grown cells after 1 week at room temperature (Bur, Cur, Rah, Mic) or after 2 weeks (Muc) as preculture suspension for time-lapse microscopy studies.

Cells were harvested from liquid precultures by sampling a 2 ml-aliquot at a culture turbidity of ca. 0.8 at 600 nm, which was centrifuged for 4 min at 8,000 rpm in a Heraeus Fresco 21 microfuge (Thermo Scientific) at room temperature. The supernatant was decanted, and the cell pellet was resuspended in 1 ml of 0.9% NaCl solution. This procedure was repeated twice more. To harvest surface-grown precultures, plates were washed with 5 ml 0.9% NaCl with a sterile 10-ml-pipette and 2 ml of this suspension was again centrifuged and washed as before. After the final resuspension, the culture turbidity was again measured, and cell suspensions were diluted with 0.9% NaCl to achieve an OD_600_ of 0.07 for Ppu or *E. coli*, 0.11 for Pve, 0.03 for Sph, Pse and Mic, 0.07 for Bur, Cur and Muc, and 0.1 for Rah. These suspensions then formed the starting point for individual cultures in time-lapse microscopy, paired mixtures (1/1 v/v ratio of the suspensions), triplicate mixtures (1/1/1 v/v) or higher orders (always maintaining equivalent volumes for each of the strains in the mixture). The culture densities were decided from various preliminary experiments (and as detailed in ^50^), such as to achieve between 30-100 founder cells per imaged area (Fig. 1e).

### Time-lapse microscopy culturing

Individual growth kinetics and behaviour, and that of pairs or higher order mixtures was measured in real-time by time-lapse microscopy imaging of surface-deposited founder cells. As support we used small solidified agarose disks (ø 10 mm and 1 mm thick) that were mounted within a stainless steel base with 42-mm ø round 0.17 mm thick coverslips on either side, as described previously ^50^. For each experiment four disks were placed, leaving ca. 1 ml air volume within the closed chamber. Media consisted of 10 g/L UltraPure™ Agarose (Invitrogen 16500-100) in 21C minimal salts medium ^54^ and carbon source(s) as detailed below. Most of the experiments with the mini-SynCom strains were conducted with soil extract as (undefined) carbon source, consisting essentially of an aqueous sterilized extract of forest soil, as described in ref ^9^, added to a concentration of ca. 100 mg C l^-1^. Alternatively, we used a mixture of 13 compounds (1 mM final concentration) thought to represent available carbon in soil ^65, 67^, or a mixture of peptone (0.5 g), trypticase (0.5 g), yeast extract (1 g) and glucose (1 g; dissolved in 1 l H_2_O containing further 0.6 g MgSO_4_·7H_2_O and 70 mg CaCl_2_·2H_2_O; called PTYG). Carbon concentrations were kept at 1 mM or 100 mg l^-1^ to avoid colonies forming multiple layers, which would have impacted the cell segmentation. The agarose (1 ml) was poured from a 60 °C-stock onto a single round 42-mm ø cover slip with a 1 mm silicone ring, loosely covered with a second cover slip until solidified. The second cover slip was then slid off horizontally and four disks with punched with a stainless steel puncher. The disks were carefully picked up with a sterile scalpel and placed in a regular four-leaf clover pattern on a new clean cover slip, after which they were exposed to the air in a sterile flow cabinet for exactly 15 minutes. Each disk was then seeded with 10 µl of individual or mixed bacterial suspension (as described above), which was distributed on the surface by placing a rectangular (1 × 1 cm) clean cover slip on top and then sliding this horizontally from the disk. The inoculated surface was then again dried to the air for exactly 15 minutes, after which a 1-mm silicon ring was added to cover the sides and the second round cover slip was placed on top, the ‘sandwich’ turned around and mounted in the closed steel chamber, cells facing down (procedure described in detail in ref. ^50^). The cover slip on the top was closed loosely with a steel screw ring not to squeeze the agarose surface, and the chamber was then transported to the microscope for imaging.

### Microscopy imaging

Growing and dividing cells on the agarose disks were imaged automatically using a Nikon ECLIPSE Ti Series inverted microscope with PFS autofocus, coupled with a Hamamatsu C11440 22CU camera and a Nikon CFI Plan Apo Lambda 100× Oil objective (1.45 numerical aperture, 1000× final magnification). The first focus was set manually and between 4 and 12 XY-positions were selected to be followed automatically through time (every 10 minutes). Positions were selected near the centre of the disks (Fig. 1a) for experimental consistency, because an oxygen gradient forms transectionally and colonies develop more profusely near the edges ^50^. Cells were imaged in phase-contrast at 50 ms exposure, mCherry at 300 ms and GFP fluorescent channels at 200 ms. The resulting images were 2048×2048 pixels (1 px corresponding to 0.065 × 0.065 µm^2^) and were saved as 16-bit.tif files. In general, we attempted to cover 24 h of growth, which was sufficient to observe exponential and stationary phases of all strains on all substrates. Despite the best of our handling practice, in some experiments the images drifted too much in XY-direction or focus was lost during the time-lapse, in which case shorter time-spans of the experiments were selected from the image series.

### Image analysis

Images were segmented in a hybrid Python pipeline named Dimalis ^68^ that combines image sharpening (BM3D ^69^), cell segmentation by Omnipose ^70^ and cell tracking by Strack ^71^. Dimalis uses the binary cell masks of the Omnipose-segmentation in phase-contrast for simultaneous segmentation of the fluorescence images at the same time point. Time-lapse series were processed by position, producing three tabular outputs (for phase-contrast, GFP and mCherry channel) with cell geometric positions, shape characteristics and fluorescence intensities. The tables were then combined into a single master table on the basis of each cell’s geometric positions, which formed the input for the cell tracking procedure (implemented in MATLAB, vs 2023, MathWorks Inc.). Cell tracking attempts to follow mother and daughter cells over multiple images across time points to build cell lineages (here taken as colonies), from which growth kinetic parameters (e.g., cell division rates, colony (lineage) area expansion rates, final productivity as final area) can be calculated. To overcome the many possible segmentation errors (e.g., missing cells, double cells as one, cell movement, blurred images, image drift), the cell tracking program reforms a bacterial cell object from the segmentation characteristics that can be restricted to or split into a minimum-maximum size (length, thickness), for which it computes the orientational overlap with cell objects on two subsequent images and retains the object with the minimal distance of the projected cell rectangular segments. The output of the cell tracking is a table with unique IDs for every cell, their attributed lineage, mother-daughter connections, cell shape characteristics and the cell’s life-time (taken as generation time).

In a second step, the sum of the cell areas in each lineage over time is calculated to estimate the colony maximum growth rate. A moving interval across 7 time points on ln-transformed summed cell areas is used to calculate slopes, of which those with an *R*^2^ above 0.95 are kept and averaged to estimate the colony’s maximum specific growth rate (µ_max_, under the assumption that the area is a proxy for the cell volume and mass). As a second measurement for the lineage growth rate we measure cell division events to calculate individual generation times, and convert these into growth rates (h^-1^) as the ratio of ln(2) divided by generation time (in h). The mean of the individual growth rates across a lineage, excluding zeros and those above 0.8 (that we assume may be the result of false cell segmentations), is then reported as the *solo* growth rate.

Next, we identify the strains on the images. Ppu and Pve are automatically assigned based on their eGFP and mCherry fluorescence, respectively, above the threshold in quantile-quantile plotting of all fluorescence values across the image. In pairs with Ppu or Pve, or in trios with Ppu and Pve, the other strain was then automatically taken as the intended partner. For other mixtures, we used a manual subroutine implemented in MATLAB that opens three phase-contrast images at early and mid exponential phase time points and stationary phase, plus a corresponding fluorescence image of eGFP and mCherry. The subroutine goes through all lineages not labeled as Ppu or Pve, placing the geometric XY-positions of the lineage founder cell on each image. Muc is then identified on the basis of its typical wider cell-cell distance at a later time point (Fig. 1b), Rah based on its resemblance to Ppu (but absence of fluorescence) and fast growth in an early time point, Cur and Mic on the basis of their typical V-cell shape in early time points (while Mic being smaller than Cur), and Bur on the basis of its slower growth than Rah and typical parallel side-by-side cells in early growth (Fig. 1b). Non-growing cells or those without one of the characteristics listed here were labelled as ‘other’ and not taken into further consideration.

### Interspecific interactions

In absence of any formality on quantification of interspecific interactions, we deployed the following concepts under the assumption that interacting cells will change growth kinetic behaviour as a function of spatial distance. This can be taken at the level of individual colonies (lineages), in which case we calculate for every colony (using the geometric XY-position of the founder cell) in circles of increasing diameter: (i) the number of founder cells of the lineages of the same kin or of the paired non-kin within the circle, and their summed cell areas at any time point. The slope of this relation is an indication for the competitive strength and direction of the interaction (Fig. 5a). Slope values with Pearson coefficient (*P* < 0.05) and linear regression coefficient (r^2^>0.2) were used to plot interaction changes over time (e.g., Fig. 5c). Slope values were further summarized across different experiments by averaging within radii of 19-45 µm, and time points within exponential phase (as in Fig. 5e). Slopes were also determined in trio mixtures for exclusive pairs within the trios (i.e., no third partner within the applied circle diameter).

In addition to interaction measurements on identified cell lineages, we deployed individual cell measurements. By placing again circles of increasing diameter around each individual cell (at each time point), we quantified changes in generation time (and its ln(2)-derived growth rate) as a function of the presence of cells of the same kin or of non-kin partners species, and compared this to growth rates of cells that are not surrounded by any other species (within the circle area). Here we report only the differences of growth rates in exclusive pairs (i.e., only kin and a single other non-kin species present within the circle), and their solo growth rates (i.e., only kin cells within the circles, as in Fig. 6).

### Spatial growth model

We use a MATLAB-embedded spatial growth model to describe microcolony formation from single cell founder cells as a function of Monod-utilization of substrate, as previously described ^50^. The model was initiated for 7 species with the same spatial positions of founder cells as in the experimental data (*n* = 8 replicates), and either mean growth rates and yields measured from surface-grown monocultures (Fig. 3a and b), or in the mini-SynComs itself (means of maximum growth rates on those lineages not surrounded by any non-kin cells within circle areas of up to 20 µm diameter, and restricted to time points between 3 and 6 hours). We further simulated the effects of changing maximum growth rates observed in paired experiments as a factor to impose on those founder cells in simulation that are closer than 15 µm from each other, or of changing biomass ratios, with the mean paired slope values of Supplementary figure 4 as a factor imposed on the per carbon substrate formed biomass yield for founder cells less than 15 µm apart. The pair-wise ratio constraint was calculated a posteriori on the total formed biomass of the mini-SynComs to estimate the species relative abundances that would best represent the measured paired competitive slopes.

## Statistical analyses

Replicates for most of the time-lapse experiments consisted of different imaged positions (FOI-technical replicates, *n* = 4–12 depending on the quality of the time-lapse imaging), and for some strain monocultures, pairs or higher order mixtures of independently started agarose patch experiments (see Supplementary figures 5 and 6 for an overview). Growth kinetic parameters among monoculture experiments (Fig. 3a) were compared by ANOVA followed by post-hoc testing of pairs (implemented in MATLAB as *anova1* and *multcompare*). Differences in strain productivities, growth rates and time to first division in pairs versus monocultures were compared in a two-sided Wilcoxon test (*ranksum*, Fig. 2). Mean net productivities in all mono-and mixed cultures were compared in a generalized linear mixed model (implemented as *fitglme,*’Distribution’,’Normal’,’Link’,’identity’,’FitMethod’,’Laplace’,’DummyVarCoding’,’effects’) testing the effect of species order (i.e., mono, pair, trio or higher) and of each strain individually (Fig. 4c). Observed paired slope values were compared to the theoretically calculated slopes from the respective measured monoculture maximum growth rates (as in Fig. 3a), according to the equation of Fig. 5a, presented as their log_2_-ratio and corresponding two-sided Wilcoxon test on individual replicate values (Fig. 5d). Differences in single cell-derived growth rates of kin and non-kin cells within the same circle radii and solo kin cells only (as in Fig. 6b) were calculated from two-sided Wilcoxon tests on individual cell values at four specific time points in exponential phase (3.4, 5, 6.6 and 8.2 h). This was used to calculate the proportion of positively or negatively deviating cell growth rates with *P* < 0.05 (as in Fig. 6b), and to define the radius at which the maximum log_2_-change in cell growth rates occurred (as in Fig. 6e). Pearson correlations and classical multi-dimensional scaling of Bray-Curtis distance dissimilarity were used to describe the outcomes of mini-SynCom simulations in comparison to the SE observations (as in Fig. 7).

## Data availability

The complete Docker version of Dimalis, including test images and user description, can be downloaded and installed from https://github.com/Helena-todd/Dimalis and https://github.com/Helena-todd/Dimalis_fluo. The additional MATLAB segmentation module and the spatially explicit agent-based growth model is downloadable with instructions from https://github.com/IsalineLucille22/Bacterial-community-growth-from-first-kinetic-principles-IBM-and-Lineages-tracking.

## Acknowledgments

The authors thank Elvire Sarton-Lohéac for her help in initial testing of the time-lapse imaging. We thank Anna Weiss and Martin Ackermann for helpful critical comments on the manucript, and all members of the NCCR for stimulating discussions during the various stages of this work.

Parts of the abstract were improved for clarity by use of large-language models and verified by human authors.

## Author contributions

T.M.T., I.G., X.R., C.M. and J.M. conceived the studies and designed experiments. T.M.T and J.M. conducted surface growth experiments and cell segmentation. H.T. and I.G. designed and implemented the imaging pipeline. I.G. developed and implemented the surface growth model. T.M.T., X.R. and J.M. analysed growth and interaction data. T.M.T. and J.M. wrote the draft manuscript. All authors provided input, verified, corrected and/or consented the final manuscript.

