## Supplementary information for "Spatially mediated interactions shape founder-cell fitness and community assembly in multi-species soil bacteria"

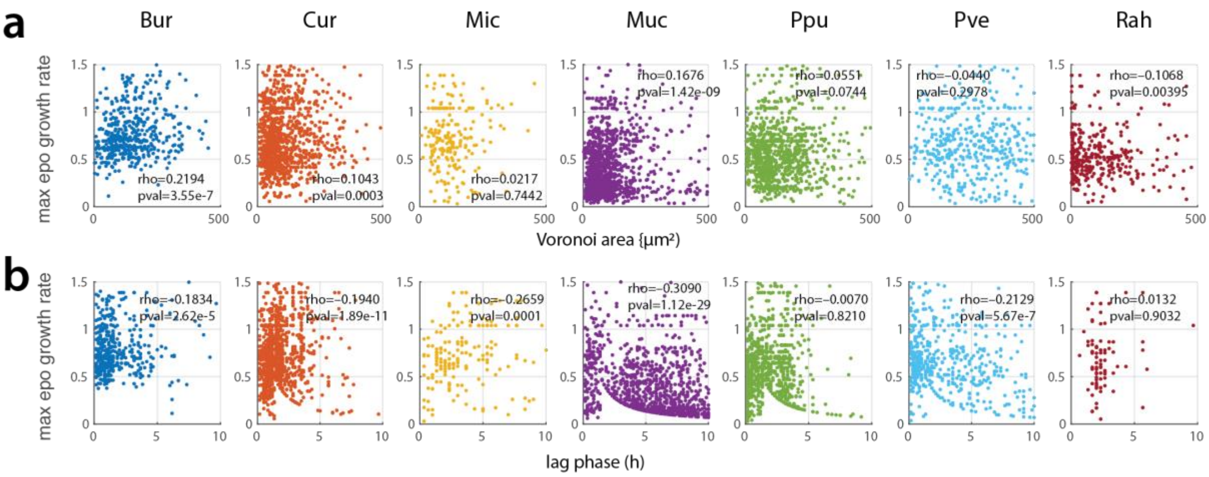


**Supplementary figure 1.** Maximum microcolony growth rate ($h^{-1}$) dependency in monocultures on (**a**) founder cell Voronoi area (in µm), or (**b**) time to first division – lag phase (h). Rho and pval correspond to the calculated Spearman correlation coefficient and its *P*-value, respectively.

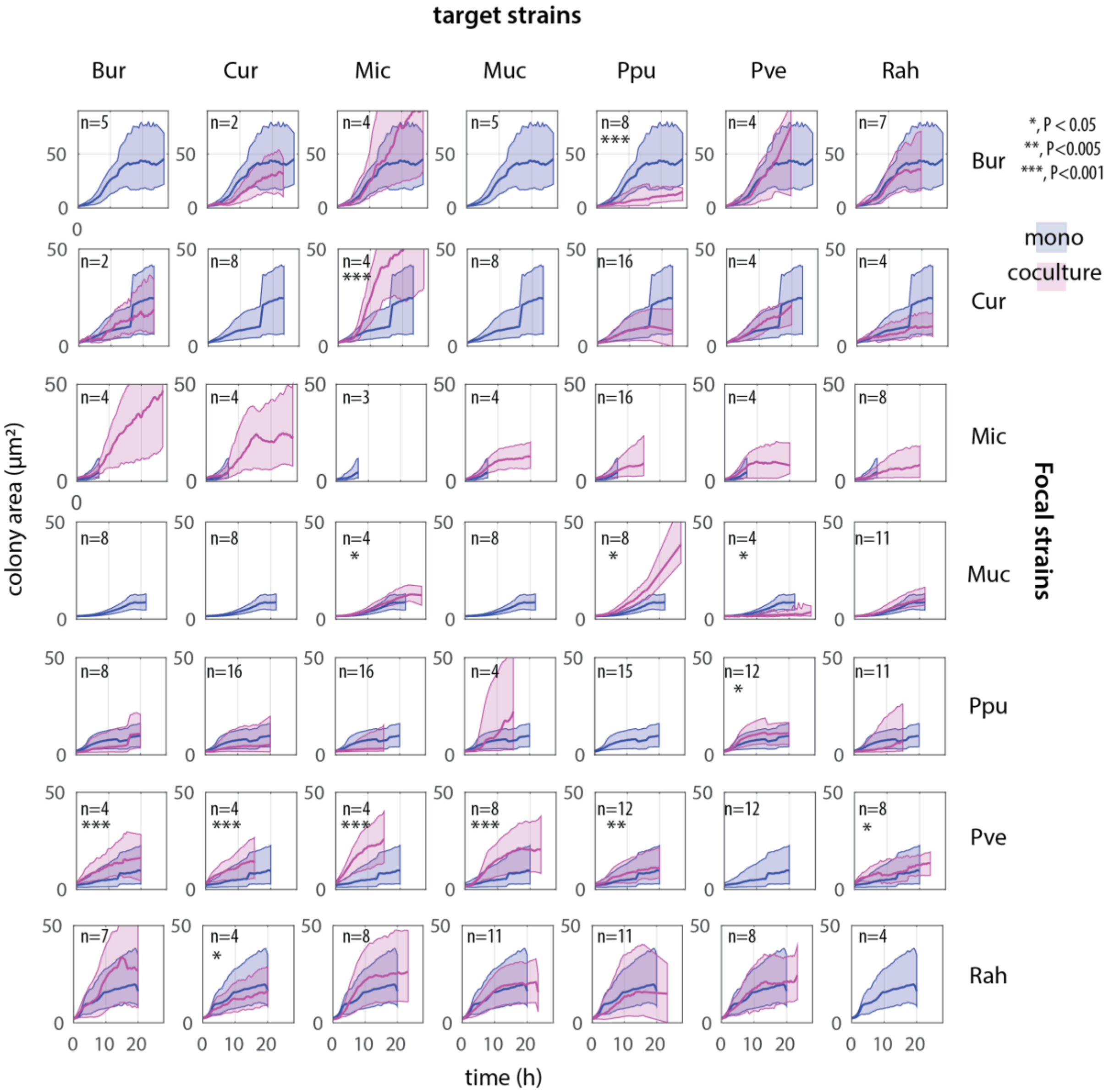


**Supplementary figure 2.** Average microcolony growth in mono- (blue hues) and paired (magenta hue) cultures of the indicated focal strains in presence of the target strains. Darker lines indicate the median and the outlines indicate the 25 and 75 th percentiles of the measured areas of all growing colonies for that strain or strain pair. n, number of replicates. *, ** and ***, thresholds for *P*-values calculated from two-sided t-test on the mean values for the averaged microcolony growth in apparent stationary phase between the mono- and paired cultures.

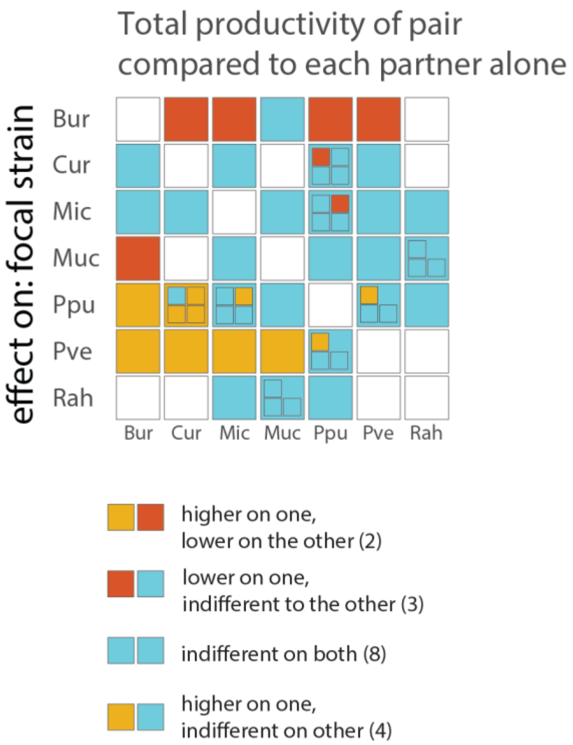


**Supplementary figure 3.** Effect of strain pairing on the total productivity. Heatmap shows change in summed productivity of the paired culture compared to either mono-culture (colour summary, full data in Supplementary figure 5b). Inset squares show individual biological replicates.

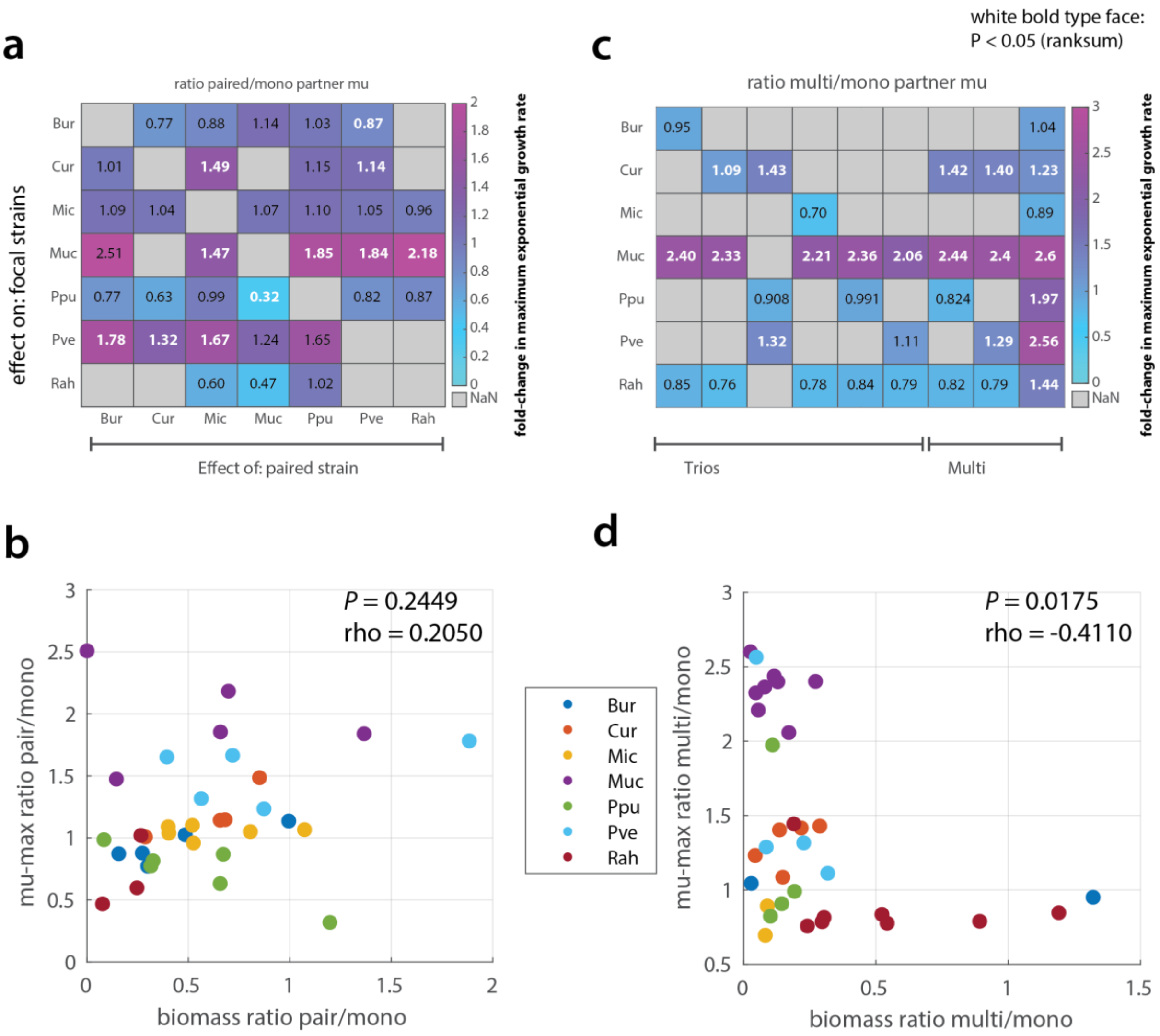


**Supplementary figure 4.** Changes of maximum specific cell lineage growth rates in species pairs, trios or higher-order mixtures. **a** Ratio of the maximum cell lineage growth rate (mu-max) in pairs and monocultures, here taken as the average of the means per imaged areas. Values in white-bold typeface are statistically significantly different (*P* < 0.05 in Wilcoxon two-sided ranksum test). **b** Correlation of the mu-max ratio per species and the final observed biomass ratio of paired compared to monoculture conditions. Dot colors correspond to the focal species as per the colorscale on the right. Each dot is an replicate (technical and/or biological replicates combined). **c** as (a), but for the ratio of mu-max in higher-order species mixtures and that in monoculture. **d** as (b) but for higher-order species mixtures.

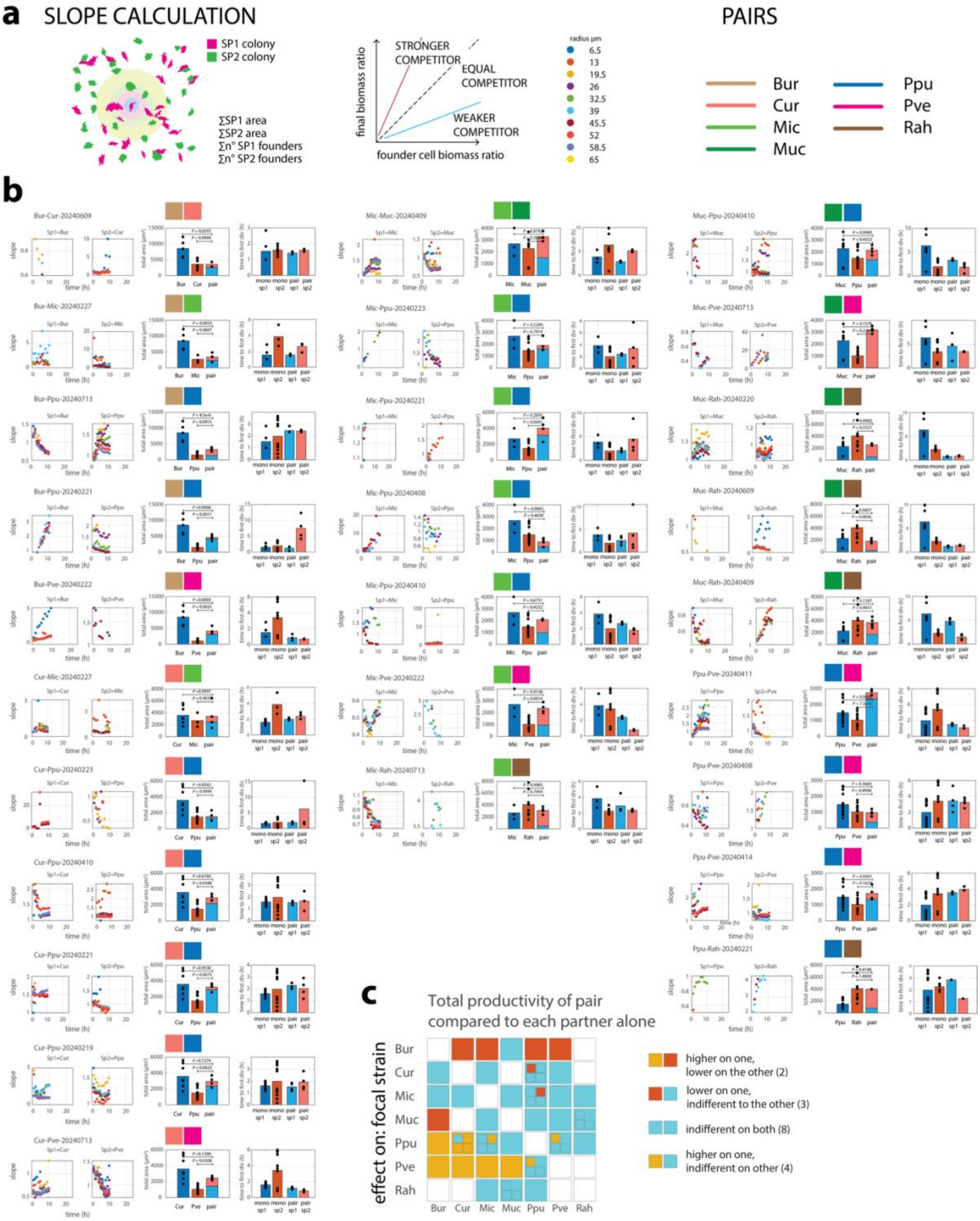


**Supplementary figure 5.** Overview of the changes in local competition index over time in all tested paired culture combinations. **a**, Principle of the calculation of the ratio between the paired species biomass at any time point and the ratio of the founder cells within circle areas of increasing diameter around every founder cell and per species. **b,** Individual plots of the change in slope across circle areas (colored dots corresponding the diameters in panel a) over time (only shown when the Pearson correlation coefficient P-value of the slope line <0.05 and the linear regression coefficient >0.2). Side plots show the mean total species biomass area per imaged area for the monocultures (blue or red, with species name below) and the coculture (as stacked bar plot of the corresponding species in light blue and light red). *P*-values from one-way Anova followed by post-hoc testing. Second side plot shows the mean time to first cell division. Color codes correspond to species combinations. **c**, Simplified plot as shown in Supplementary figure 3 for the paired versus monoculture productivities, deduced from the individual side plots of productivity.

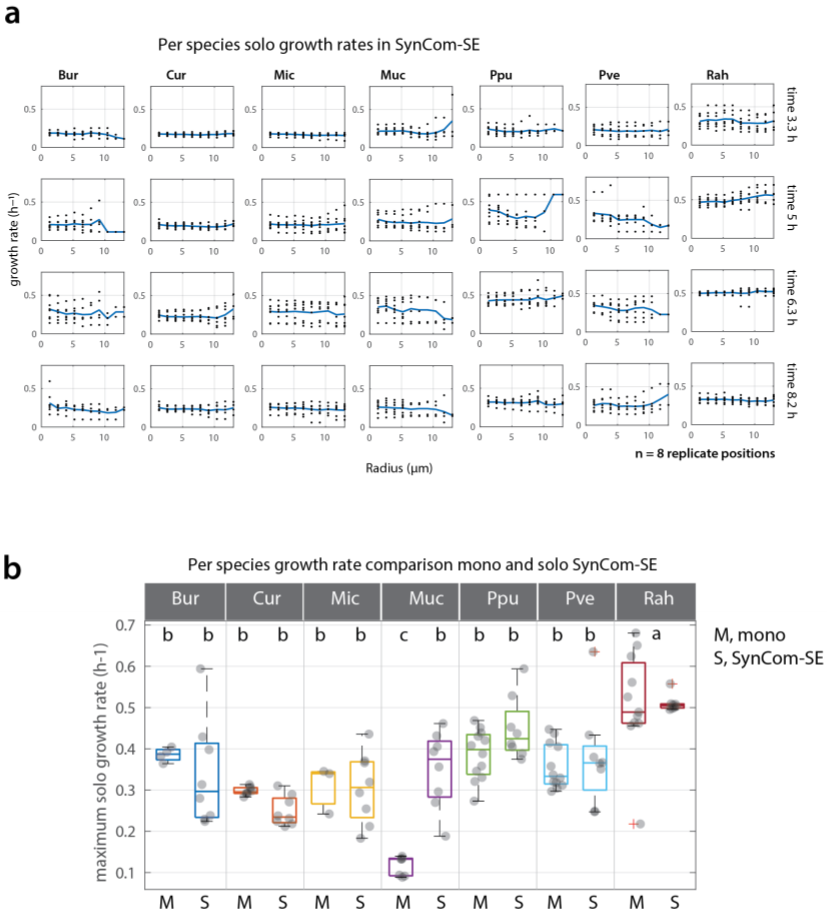


**Supplementary figure 6.** Per species maximum solo growth rates. **a** Growth rates converted from generation times of individual cells of the indicated species at four different timepoints growing within the seven-member community on soil extract (SE), sampled in areas without any other species neighbours within a radial distance of 15 µm (i.e., "solo" growth rates). Dots are means of individual cells within the indicated radius for eight technical replicates (i.e., imaged areas). Thick blue lines connect the means. The highest average over all radii of the means for any of the time points is taken as the maximum solo growth rate of the species. **b** Comparison of per species maximum solo growth rates in monoculture and within the seven-member community growing on SE (S, SynCom-SE). Dots are the highest average over all radii (as in **a**) but for the individual replicates. Representation is a regular box-plot with 25th, median and 75th percentiles. Letters (a, b, c) correspond to significantly different groups in one-way ANOVA followed by post-hoc testing.

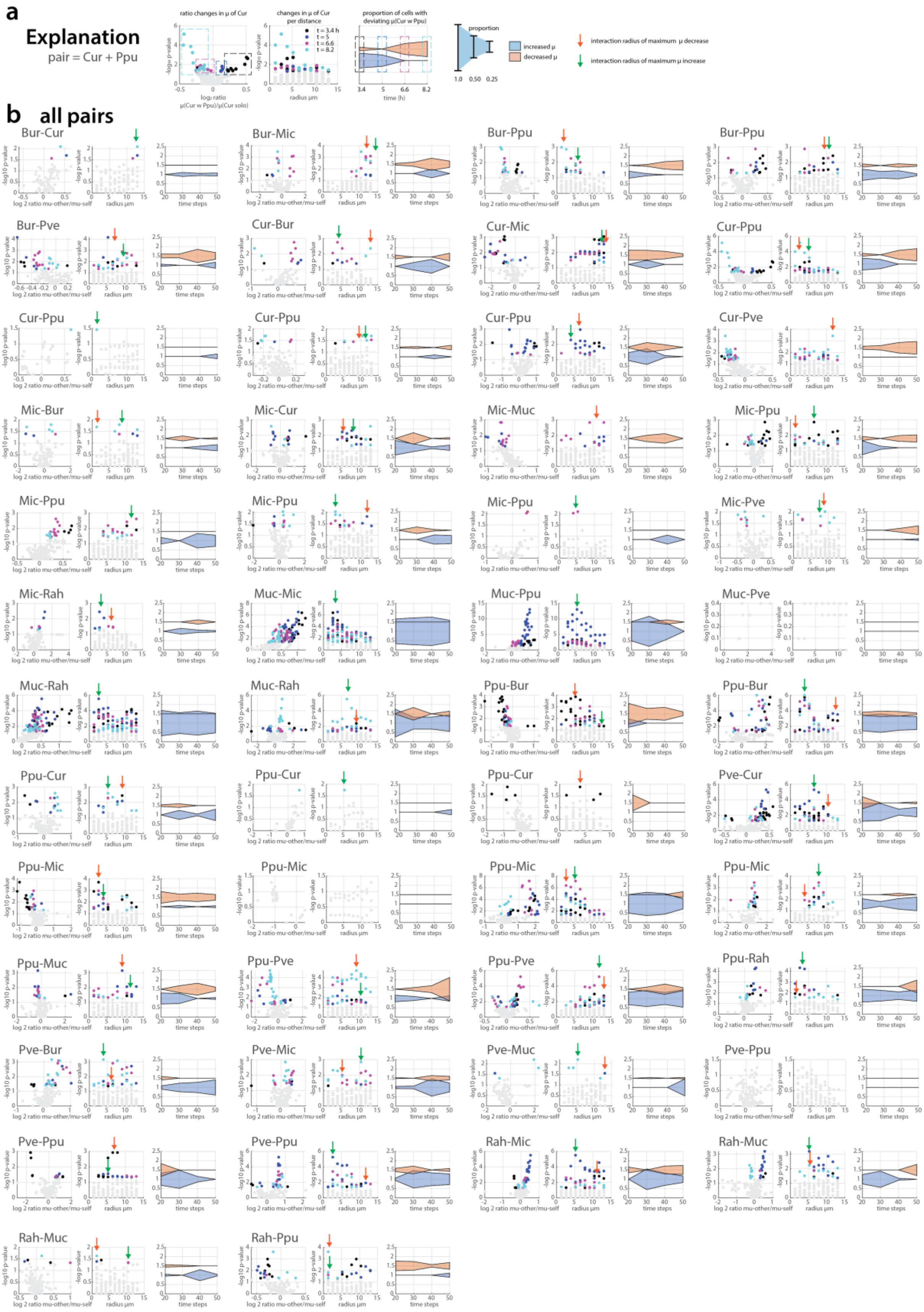


**Supplementary figure 7.** Individual cell-based interaction plots in paired species combinations (background data for Fig. 6). **a** Rehearsal of the explanation of the concept of the paired species individual cell measurements. **b** Plots for all indicated species combinations (some in multiple independent replicated patch incubations) show the $-\log_{10}$ *P*-value of the Wilcoxon test of the measured growth rate of cells in presence of a non-kin partner compared to that of only kin-cells, within circles of increasing radius, as a function of the $\log_2$ ratio of those growth rates. Grey dots have a *P*-value > 0.05, colored dots are lower. Color corresponds to the measured time point (black is 20 time steps or 3.3 h; 30 = 5 h, 40 = 6.7 and 50= 8.2 h). Next plot shows the $-\log_{10}$ *P*-value as a function of the circle radius (in µm), and finally the derived proportion of cells with statistically significantly higher (blue) or lower (salmon) growth rates over time (these plots are reported in Figure 6d). 6 time steps correspond to 1 h. Green and red arrows signal the radius of the maximum derived positive or negative occurrence (among the significant points).

**Supplementary table 1. Percentage of number of focal strain cells of the (exclusive) pair within circle radius 15 µm compared to all cells of focal strain (at time = 5 h within the mini-SynCom grown on SE).**

| focal strain | partner strain | pos1 | pos2 | pos3 | pos4 | pos5 | pos6 | pos7 | pos8 | mean |
|---|---|---|---|---|---|---|---|---|---|---|
| Bur | Cur | 0.00% | 0.00% | 0.00% | 0.00% | 0.00% | 0.00% | 0.00% | 7.69% | 0.96% |
| Bur | Mic | 33.33% | 0.00% | 50.00% | 0.00% | 4.55% | 3.45% | 0.00% | 7.69% | 12.38% |
| Bur | Muc | 0.00% | 0.00% | 0.00% | 0.00% | 0.00% | 0.00% | 0.00% | 0.00% | 0.00% |
| Bur | Ppu | 0.00% | 0.00% | 0.00% | 0.00% | 0.00% | 0.00% | 0.00% | 38.46% | 4.81% |
| Bur | Pve | 0.00% | 0.00% | 0.00% | 0.00% | 0.00% | 0.00% | 0.00% | 0.00% | 0.00% |
| Bur | Rah | 8.33% | 0.00% | 0.00% | 0.00% | 13.64% | 6.90% | 0.00% | 0.00% | 3.61% |
| Cur | Bur | 7.41% | 0.00% | 0.00% | 27.59% | 5.71% | 0.00% | 0.00% | 1.28% | 5.25% |
| Cur | Mic | 0.00% | 0.00% | 0.00% | 3.45% | 0.00% | 0.00% | 9.68% | 12.82% | 3.24% |
| Cur | Muc | 0.00% | 0.00% | 0.00% | 10.34% | 0.00% | 0.00% | 3.23% | 0.00% | 1.70% |
| Cur | Ppu | 0.00% | 0.00% | 0.00% | 0.00% | 0.00% | 0.00% | 0.00% | 6.41% | 0.80% |
| Cur | Pve | 0.00% | 0.00% | 0.00% | 0.00% | 17.14% | 4.00% | 0.00% | 0.00% | 2.64% |
| Cur | Rah | 14.81% | 5.00% | 6.25% | 0.00% | 2.86% | 16.00% | 6.45% | 0.00% | 6.42% |
| Mic | Bur | 5.77% | 0.00% | 0.00% | 0.00% | 0.00% | 9.09% | 2.04% | 1.27% | 2.27% |
| Mic | Cur | 0.00% | 0.00% | 5.75% | 0.00% | 0.00% | 0.00% | 3.06% | 18.99% | 3.47% |
| Mic | Muc | 0.00% | 0.00% | 9.20% | 0.00% | 0.00% | 2.27% | 4.08% | 0.00% | 1.94% |
| Mic | Ppu | 0.00% | 6.00% | 10.34% | 0.00% | 8.70% | 2.27% | 0.00% | 1.27% | 3.57% |
| Mic | Pve | 5.77% | 0.00% | 3.45% | 22.92% | 0.00% | 2.27% | 0.00% | 0.00% | 4.30% |
| Mic | Rah | 13.46% | 0.00% | 1.15% | 0.00% | 17.39% | 2.27% | 6.12% | 1.27% | 5.21% |
| Muc | Bur | 0.00% | 5.88% | 0.00% | 0.00% | 0.00% | 0.00% | 0.00% | 3.57% | 1.18% |
| Muc | Cur | 0.00% | 0.00% | 0.00% | 12.50% | 0.00% | 0.00% | 0.00% | 7.14% | 2.46% |
| Muc | Mic | 0.00% | 0.00% | 30.00% | 0.00% | 0.00% | 0.00% | 20.00% | 0.00% | 6.25% |
| Muc | Ppu | 0.00% | 0.00% | 0.00% | 0.00% | 28.57% | 0.00% | 20.00% | 0.00% | 6.07% |
| Muc | Pve | 0.00% | 0.00% | 0.00% | 0.00% | 0.00% | 0.00% | 0.00% | 0.00% | 0.00% |
| Muc | Rah | 0.00% | 0.00% | 0.00% | 12.50% | 0.00% | 0.00% | 0.00% | 0.00% | 1.56% |
| Ppu | Bur | 0.00% | 0.00% | 9.09% | 0.00% | 0.00% | 0.00% | 0.00% | 30.00% | 4.89% |
| Ppu | Cur | 0.00% | 0.00% | 0.00% | 0.00% | 0.00% | 0.00% | 0.00% | 0.00% | 0.00% |
| Ppu | Mic | 0.00% | 0.00% | 27.27% | 0.00% | 30.77% | 0.00% | 4.17% | 40.00% | 12.78% |
| Ppu | Muc | 0.00% | 0.00% | 0.00% | 0.00% | 38.46% | 0.00% | 0.00% | 0.00% | 4.81% |
| Ppu | Pve | 0.00% | 0.00% | 0.00% | 0.00% | 0.00% | 0.00% | 0.00% | 0.00% | 0.00% |
| Ppu | Rah | 0.00% | 0.00% | 0.00% | 0.00% | 7.69% | 7.50% | 0.00% | 0.00% | 1.90% |
| Pve | Bur | 0.00% | 0.00% | 0.00% | 0.00% | 27.27% | 0.00% | 0.00% | 0.00% | 3.41% |
| Pve | Cur | 0.00% | 50.00% | 2.13% | 0.00% | 9.09% | 6.67% | 0.00% | 0.00% | 8.49% |
| Pve | Mic | 21.43% | 0.00% | 8.51% | 55.00% | 0.00% | 20.00% | 0.00% | 0.00% | 13.12% |
| Pve | Muc | 0.00% | 0.00% | 0.00% | 0.00% | 0.00% | 0.00% | 0.00% | 0.00% | 0.00% |
| Pve | Ppu | 0.00% | 0.00% | 0.00% | 0.00% | 0.00% | 0.00% | 16.67% | 0.00% | 2.08% |
| Pve | Rah | 0.00% | 0.00% | 17.02% | 10.00% | 0.00% | 0.00% | 0.00% | 0.00% | 3.38% |
| Rah | Bur | 8.16% | 0.00% | 0.00% | 0.00% | 3.37% | 4.17% | 0.00% | 0.00% | 1.96% |
| Rah | Cur | 9.18% | 15.79% | 1.64% | 26.47% | 0.00% | 13.54% | 3.88% | 0.00% | 8.81% |
| Rah | Mic | 26.53% | 6.58% | 0.00% | 0.00% | 23.60% | 4.17% | 24.27% | 0.00% | 10.64% |

| | | | | | | | | | | |
|---|---|---|---|---|---|---|---|---|---|---|
| Rah | Muc | 0.00% | 0.00% | 0.00% | 5.88% | 0.00% | 0.00% | 0.00% | 0.00% | 0.74% |
| Rah | Ppu | 0.00% | 0.00% | 37.70% | 0.00% | 0.00% | 0.00% | 1.94% | 0.00% | 4.96% |
| Rah | Pve | 0.00% | 0.00% | 4.92% | 8.82% | 0.00% | 0.00% | 0.00% | 0.00% | 1.72% |

**Supplementary table 2. Total percentage of cells of focal strain engaged in exclusive pairs at time 5 h within the mini-SynCom grown on SE.**

| strain | pos1 | pos2 | pos3 | pos4 | pos5 | pos6 | pos7 | pos8 | mean |
|---|---|---|---|---|---|---|---|---|---|
| Bur | 41.67% | 0.00% | 50.00% | 0.00% | 18.18% | 10.34% | 0.00% | 53.85% | 21.75% |
| Cur | 22.22% | 5.00% | 6.25% | 41.38% | 25.71% | 20.00% | 19.35% | 20.51% | 20.05% |
| Mic | 25.00% | 6.00% | 29.89% | 22.92% | 26.09% | 18.18% | 15.31% | 22.78% | 20.77% |
| Muc | 0.00% | 5.88% | 30.00% | 25.00% | 28.57% | 0.00% | 40.00% | 10.71% | 17.52% |
| Ppu | 0.00% | 0.00% | 36.36% | 0.00% | 76.92% | 7.50% | 4.17% | 70.00% | 24.37% |
| Pve | 21.43% | 50.00% | 27.66% | 65.00% | 36.36% | 26.67% | 16.67% | 0.00% | 30.47% |
| Rah | 43.88% | 22.37% | 44.26% | 41.18% | 26.97% | 21.88% | 30.10% | 0.00% | 28.83% |
